# HIV broadly neutralizing antibody precursor maturation induced by mRNA vaccination in nonhuman primates

**DOI:** 10.64898/2026.11.10.765717

**Authors:** Ivy Phung, Brian L. Freeman, Alessia Liguori, Amber Seymour, Zhenfei Xie, Swastik Phulera, Oleksandr Kalyuzhniy, Patrick J. Madden, Ester Marina-Zárate, Jordan R. Willis, Christopher A. Cottrell, Krystal M. Ma, Sabyasachi Baboo, Claudia T. Flynn, Carolyne Kifude, Danny Lu, Nicole Phelps, Diana Goodwin, Yumiko Adachi, Ryan Tingle, Erik Georgeson, Jenny Tran, Anton Kalyuzhniy, Xiaozhen Hu, Vanessa R. Lewis, Julia T. Ngo, Xiaotie Liu, Jolene K. Diedrich, Usha Nair, Sunny Himansu, Torben Schiffner, Paolo Lusso, James C. Paulson, John R. Yates, Diane G. Carnathan, Dennis R. Burton, Darrell J. Irvine, Guido Silvestri, Andrew B. Ward, Gabriel Ozorowski, Facundo D. Batista, Parham Ramezani-Rad, Jon M. Steichen, Shane Crotty, William R. Schief

**Author notes:** These authors contributed equally to this work (Ivy Phung, Brian L. Freeman). Co-corresponding authors: William R. Schief, The Scripps Research Institute 10550 N Torrey Pines Rd, La Jolla, CA 92037, Shane Crotty La Jolla Institute for Immunology 9420 Athena Cir, La Jolla, CA 92037, Jon M. Steichen, The Scripps Research Institute 10550 N Torrey Pines Rd, La Jolla, CA 92037, Parham Ramezani-Rad La Jolla Institute for Immunology 9420 Athena Cir, La Jolla, CA 92037, Facundo D. Batista, Ragon Institute of Mass General Brigham 600 Main St, Cambridge, MA 02139.

## Abstract

The induction of broadly neutralizing antibodies (bnAbs) is a principal goal of prophylactic HIV vaccines. For many bnAbs, priming ultrarare precursors from the naïve B cell repertoire is the first major immunological hurdle. Previous studies have shown that the germline-targeting (GT) priming immunogen N332-GT5 delivered as adjuvanted protein to non-human primates (NHPs) activates bnAb precursor B cells for the human bnAb BG18 targeting the V3-glycan supersite on the HIV envelope (Env) glycoprotein. Here, we engineered N332-GT6 to improve binding affinity to diverse BG18-class precursor naive B cells by 9-fold. mRNA-LNP vaccines expressing N332-GT5 or N332-GT6 as membrane-bound (gp151) trimers were evaluated for their ability to prime BG18-class precursor B cells in NHPs across three related studies. Although N332-GT5 gp151 mRNA-LNP priming induced BG18-class precursor B cells in most animals after two vaccinations, the higher-affinity N332-GT6 gp151 induced responses more consistently after only one immunization. Sequential vaccination with two heterologous mRNA-LNP booster immunizations drove affinity maturation and induced BG18-class memory B cells capable of binding wildtype HIV Env. Together, these findings demonstrate that mRNA immunization can achieve GT priming, boosting, and early maturation of ultrarare bnAb-class B cells in NHPs.

## Introduction

Prophylactic vaccines are among the most effective preventative modalities in modern medicine. HIV vaccine development has remained a major immunological challenge for over four decades because of extensive genetic diversity of the HIV envelope glycoprotein (Env), as well as HIV Env structural features that limit the immunogenicity of conserved neutralizing epitopes^1^. However, the discovery of broadly neutralizing antibodies (bnAbs) in people living with HIV that potently neutralize diverse HIV-1 isolates has aided research efforts towards an antibody-based vaccine^2^. bnAbs typically arise through years of antibody-virus coevolution, acquiring extensive somatic hypermutation (SHM) and high affinity, whereas their naïve B cell precursors are generally unable to bind wildtype HIV Env glycoproteins^3^. Compounding this challenge, HIV bnAb-precursor B cells can be exceedingly rare and require unique structural components to facilitate antigen-binding^4^. Germline-targeting (GT) vaccine design aims to elicit bnAbs by priming rare bnAb-precursor B cells and guiding their affinity maturation through sequential immunization with increasingly native-like Env booster immunogens^1,4–11^.

Transgenic mouse models that utilized knock-in human B cell receptors (BCR), recombining human immunoglobulin genes^12,13^, or adoptive B cell transfer experiments with physiological precursor frequencies^14^, as well as studies in NHPs, have demonstrated successful priming and competitive fitness of bnAb-class B cells across multiple Env epitopes. Examples include eOD-GT8 60mer (VRC01-class)^9,13–17^, 10E8-GT (MPER)^10,18^, Apex-GT (V2-apex)^11,19,20^, N332-GT (BG18/PGT121-class)^4,21–24^, and combination priming with up to four different bnAb classes^26,27^. Subsequent “shepherding” with more native-like immunogens has also been validated in multiple animal models^13,21,23,25,28^. Clinical proof-of-concept was established in a first-in-human GT priming study, where rare VRC01-class precursor B cells were primed in 97% of vaccine recipients^29^. VRC01-class antibodies were well suited as a first GT vaccine design because they rely primarily on germline-encoded HCDR2 interactions for Env binding, an interaction mode that is relatively unusual but technically more tractable for GT than more conventional HCDR3-dominant antigen binding^30,31^. HCDR3-dominant bnAbs are more common and represent a bigger challenge for GT vaccine design^4,10,24^.

Previous studies have characterized N332-GT5, a novel HCDR3-dominant GT immunogen designed to prime rare B cells targeting the V3-glycan/N332 supersite with features of the HIV bnAb BG18^4,24,32^. BG18-class precursor frequencies are exceptionally rare in humans (∼1 in 50 million)^4^ and estimated to be as much as 8-fold lower in NHPs^24,33^. Despite this, priming of BG18-class precursor B cells was achieved in NHP immunization studies using with N332-GT5 protein with SMNP adjuvant, with BG18-class B cell responses observed in almost all animals^24,33,34^. These findings supported the feasibility of priming rare HCDR3-dominant B cell precursors in humans. Recently, sequential protein immunization beginning with N332-GT5 achieved bnAb maturation in memory B cells and serum by priming and shepherding BG18-class B cells in NHPs^28^.

Recombinant proteins such as N332-GT5 have been studied extensively for HIV vaccine development. However, manufacturing protein subunit vaccines for human use can be costly and time-consuming^35^. Recombinant HIV Env trimer proteins also come with immunogenicity caveats: first, Env alone is poorly immunogenic and requires potent and safe adjuvants to enhance immune responses; and second, soluble Env exposes the immunodominant non-neutralizing trimer base^36^. Several strategies to address these challenges have been proposed, which include next-generation adjuvants^34,37,38^, modulating antigen delivery and retention^39–41^, and protein engineering to mask off-target epitopes^33,42–44^.

Using mRNA-LNP as an HIV vaccine platform is a promising approach for expression of immunogens in a more native context, such as membrane-bound Env trimers that can increase responses to subdominant epitopes in NHPs^7,11,45^ and humans^46^. GMP-grade mRNA constructs are significantly more cost-effective and easier to manufacture than GMP proteins. mRNA-based vaccines have demonstrated robust efficacy in rigorous clinical trials. During the SARS-CoV-2 pandemic, mRNA vaccines proved highly effective at preventing severe COVID-19, with extensive safety data from billions of doses administered worldwide^47,48^. mRNA platforms have also shown promise for use in GT-vaccine strategies for HIV in humans^49^. Here, we examine the potential of mRNA-LNP vaccine designs to prime and boost ultrarare BG18-class precursors in a series of three NHP studies.

## Results

### Design of an N332-GT6 immunogen targeting the HIV V3-glycan supersite

Although N332-GT5 has demonstrated very encouraging results in mice and NHPs, several properties could potentially be improved. For example, priming of BG18-class precursors in NHPs by bolus administration of N332-GT5 was first detected in the circulating memory B cell (B_Mem_) pool after a booster immunization^33^. The number of unique BG18-class B cell lineages detected after either escalating dose (ED) or bolus priming was 0-12 per animal, with a median of 2 to 3.5 lineages^33^. A goal of GT priming is to engage multiple BG18-class B cell lineages per subject to maximize the likelihood of subsequent successful maturation of BG18-class bnAbs^29^. An N332-GT with improved affinity may prime a more diverse set of precursors and sustain a greater number of lineages. Engaging more BG18-class B cell lineages could increase the likelihood that sequential immunization with a series of heterologous boosters can elicit affinity matured bnAbs. Additionally, N332-GT5 protein has a low yield when expressed by transient transfection in 293F cells, creating technical challenges for vaccine studies (though expression yield from stable cell lines in was quite good). Improving the expression yield of N332-GT protein could have two benefits: 1) protein immunogens and reagents could be produced at lower cost and effort and 2) mRNA immunogens might deliver protein at higher in vivo expression levels and therefore have higher immunogenicity.

Because mammalian display-directed evolution methods were successful at creating N332-GT5, we used similar methods to improve the affinity of N332-GT5 for diverse BG18-class precursors. The design goal was to identify mutations that would enhance binding to BG18-class precursors with diverse HCDR3 junctions. We assembled a collection of BG18-class precursors identified in a next-generation sequencing (NGS) dataset of human antibody HCs^4,50^. One focus was the position two amino acids N-terminal to the D gene. This position is nearly always Glu in BG18-class antibodies elicited in NHPs^24^, but Glu is infrequent (<5%) at this position in the naïve human repertoire (**Supplementary Fig. 1a**). To increase the number of potential BG18-class precursors activated during priming, we sought to increase affinity for BG18-class precursors lacking a Glu at this position, with the rationale that the Glu could be selected by subsequent heterologous boosters. Accordingly, several antibodies used as mammalian display selection reagents had amino acids other than Glu at this position (**Supplementary Fig. 1b**). In addition to these NGS-derived BG18-class precursors, we used inferred-germline BG18 paired with a variety of different light chains (LCs), with the goal of improving affinity for precursors with diverse light chain V genes beyond the V_L_3-25 gene utilized by BG18.

Starting from N332-GT5, a random mutagenesis library was first screened that covered the majority of the BG18 epitope (residues 114 to 366 in HxB2 numbering) including the V1 and V3 loops (**Fig. 1a**). This identified two mutations (N302I and K155M). Next, a structure-guided combinatorial library identified mutations E153Q and T303S that, combined with N302I and K155M, generated 25mutB. A second structure-guided library identified four additional mutations (G389L, D412E, V442P, and R444N); these were added to 25mutB to generate 29mutA, which had 5-fold higher geometric mean affinity for BG18-class precursors compared to N332-GT5 (**Fig. 1a-b**). Next, two saturation mutagenesis libraries were assembled targeting segments spanning residues 120 to 212 and 290 to 446. The most enriched mutations were incorporated into new combinatorial libraries that were screened, and 11 additional mutations were selected from those screens. When these mutations were incorporated into 29mutA, the resulting trimer, termed N332-GT6, had 19 mutations relative to N332-GT5 (**Fig. 1a**) and 38 GT mutations relative to the stabilized native-like trimer BG505 MD39. N332-GT6, like N332-GT5, also contained MD39 stabilization mutations, including SOSIP mutations and other mutations^23^.

**Fig. 1:**
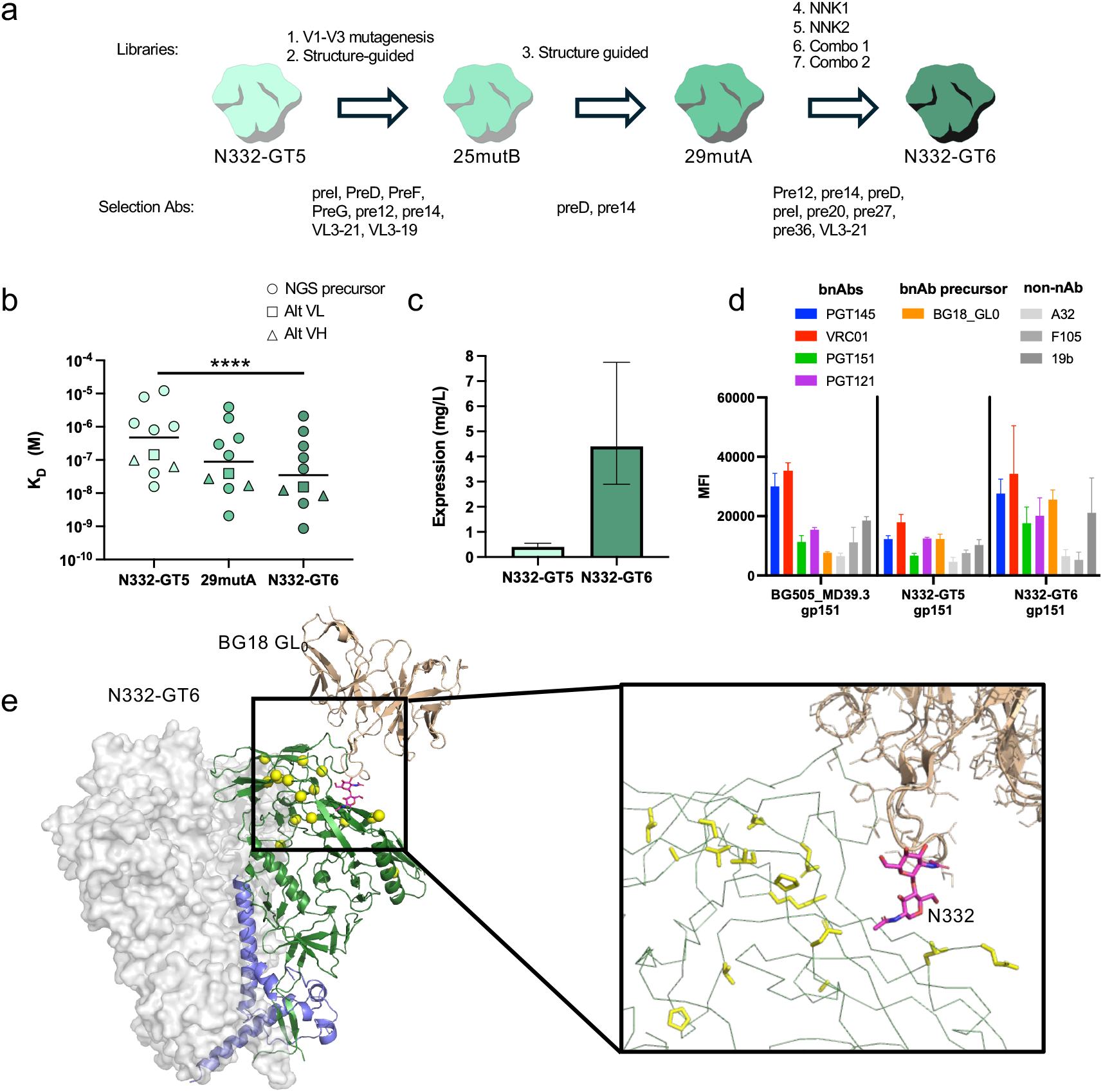
N332-GT6 design. **a,** Schematic for designing N332-GT6 by mammalian display directed evolution. **b,** Binding affinity (*K*_D_) measured by SPR for a panel of BG18 type I precursors derived from NGS search (NGS precursors), BG18 iGL with alternate VL gene (Alt VL) or VH gene (Alt VH). Statistical significance was determined using a paired *t* test on log-transformed K_D_. **c,** Expression yield for purified N332-GT6 compared to N332-GT5. **d,** Antigenic profile of membrane-bound N332-GT6 gp151 expressed on 293F cells. **e,** Structure of unliganded N332-GT6 with BG18 GL_0_ modeled based on alignment to PDB 6DFH. gp120 from one protomer shown as green cartoon with N332-GT6 mutations shown in yellow balls or sticks (inset) and the N332 glycan shown as raspberry spheres. gp41 protomer is shown in purple cartoon and BG18 GL_0_ is shown as brown cartoon. The other two protomers are shown as gray surface.

N332-GT6 had a 9-fold improved geometric mean *K*_D_ for diverse BG18-class precursors compared to N332-GT5 (*P* < 0.0001, **Fig. 1b, Supplementary Fig. 1c**). N332-GT6 also showed 9-fold improvement over N332-GT5 in geometric mean *K*_D_ for BG18-class precursors isolated after priming NHPs with N332-GT5 (*P* < 0.0001, **Supplementary Fig. 1d**) and a 36-fold improvement over N332-GT5 in geometric mean *K*_D_ for naïve precursors from BG18^gl^ HC KI mice (*P* < 0.0001, **Supplementary Fig. 1d**). Together, these data indicated that N332-GT6 has improved affinity across diverse BG18-class precursors.

An additional benefit of the mammalian display selection strategy was that soluble N332-GT6 had an 11-fold higher expression yield compared to N332-GT5 (**Fig. 1c**). Transient transfection of N332-GT6 in 293T cells as a membrane-bound gp151 Env trimer (designed as per Willis et al.^20^ and Xie et al.^21^, see Methods) resulted in expression levels ∼2-fold higher than membrane-bound N332-GT5 gp151 with an improved antigenic profile comparable to BG505 MD39.3 gp151, but with a strong gain in binding to inferred-germline (iGL) precursor antibody BG18 iGL_0_ (**Fig. 1d, Supplementary Fig. 2**). The N332-GT6 trimer was thermostable, and structural analysis of unliganded N332-GT6 showed no substantial structural changes compared to N332-GT5 (**Supplementary Fig. 1e, f, Supplementary Fig. 3**). Modeling BG18 iGL_0_ at the interface indicated that none of the GT mutations make direct contacts to BG18-class precursors, suggesting that indirect structural effects drive the enhanced affinity for BG18-class precursors (**Fig. 1e**). The soluble N332-GT6 trimer also showed a favorable antigenic profile, with substantial binding to bnAbs and minimal binding to non-nAbs (**Supplementary Fig. 1g**), and a glycan occupancy and glyocoform profile typical of a native-like Env trimer (**Supplementary Fig. 1h**).

### Evaluation of N332-GT5 and N332-GT6 mRNA-LNP in mouse models

N332-GT5 and N332-GT6 protein immunogens were compared in mouse models. Adoptive transfer experiments were performed using two newly generated humanized mouse models with NGS-derived BG18-class precursors (Pre5 and Pre7 from Steichen et al.^4^) knocked in. Pre5 and Pre7 naïve B cells (CD45.2^+^) were adoptively transferred to establish precursor frequencies at different levels including low (∼3 in 10^6^ for Pre5 and ∼2 in 10^6^ for Pre7), intermediate (12 in 10^6^ and 9 in 10^6^, respectively), and high (32 in 10^6^ and 25 in 10^6^, respectively) (**Supplementary Fig. 4**). Adoptively transferred mice were immunized with N332-GT5 and N332-GT6 protein immunogens formulated with Sigma adjuvant (**Supplementary Fig. 5a**). After these protein immunizations, BG18-class precursor priming was only detected sporadically and at high precursor frequency (**Supplementary Fig. 5b-f**).

To assess immunogenicity of N332-GT5 and N332-GT6 as mRNA-LNPs, adoptively transferred mice received intramuscular (IM) immunization with membrane-bound N332-GT5, N332-GT6.2 (without glycosylation sites at positions 241 and 289), or N332-GT6.3 (with glycosylation sites at 241 and 289, filling the “glycan hole” of BG505). Inguinal, popliteal and iliac draining lymph node samples were assayed by flow cytometry at day 13 (**Supplementary Fig. 6a**). All three mRNA-LNP constructs elicited germinal center B cell (B_GC_) responses (**Supplementary Fig. 6c-d**). At the lowest Pre5 precursor frequency, Pre5-derived B_GC_ amounted to ∼10% of B_GC_ following N332-GT6.2 immunization but were barely detectable after N332-GT5 immunization (N332-GT6.2 vs N332-GT5, *P* < 0.05; **Supplementary Fig. 6d-e**). In addition, both N332-GT6.2 and N332-GT6.3 mRNA constructs induced a trend towards a higher proportion of N332-GT-binding Pre5 B_GC_ compared with N332-GT5 mRNA under this condition (**Supplementary Fig. 6f**). Taken together, these results indicate that N332-GT5, N332-GT6.2 and N332-GT6.3 mRNA constructs can prime BG18-class precursors. N332-GT6.2 and N332-GT6.3 mRNA constructs showed greater recruitment of antigen-binding precursor B cells under some conditions in these humanized mouse models. Hereafter, N332-GT6 mRNA refers to the N332-GT6.3 construct, which contains glycosylation sites at positions 241 and 289.

### N332-GT6 mRNA primes higher frequencies of BG18-class B cells than N332-GT5 mRNA in NHPs

To assess whether N332-GT5 or N332-GT6 mRNA-LNP can prime BG18-class responses in NHPs, we performed a head-to-head comparison study of membrane-bound N332-GT6 mRNA, membrane-bound N332-GT5 mRNA, and soluble N332-GT5 gp140 protein (NHP Study 1). Four groups of six NHPs each were immunized (**Fig. 2a**). Group 1 animals were primed with a previously validated^24,33^ ED regimen of soluble N332-GT5 protein adjuvanted with SMNP and a homologous bolus booster immunization at week 10. This immunization group served as a benchmark comparison. Group 2 animals received a priming immunization of membrane-bound N332-GT5 mRNA with two homologous booster immunizations at weeks 8 and 16. Group 3 animals received the same regimen as Group 2, but with N332-GT6 mRNA. Group 4 animals received a priming immunization of N332-GT5 mRNA with a short-dosing regimen at week 2 and 10 (**Fig. 2a**). The short-dosing mRNA regimen tested whether introducing an earlier second full dose of mRNA-LNP improved immune responses, inspired by a 2-shot ED protein regimen tested in mice^51^.

**Fig. 2:**
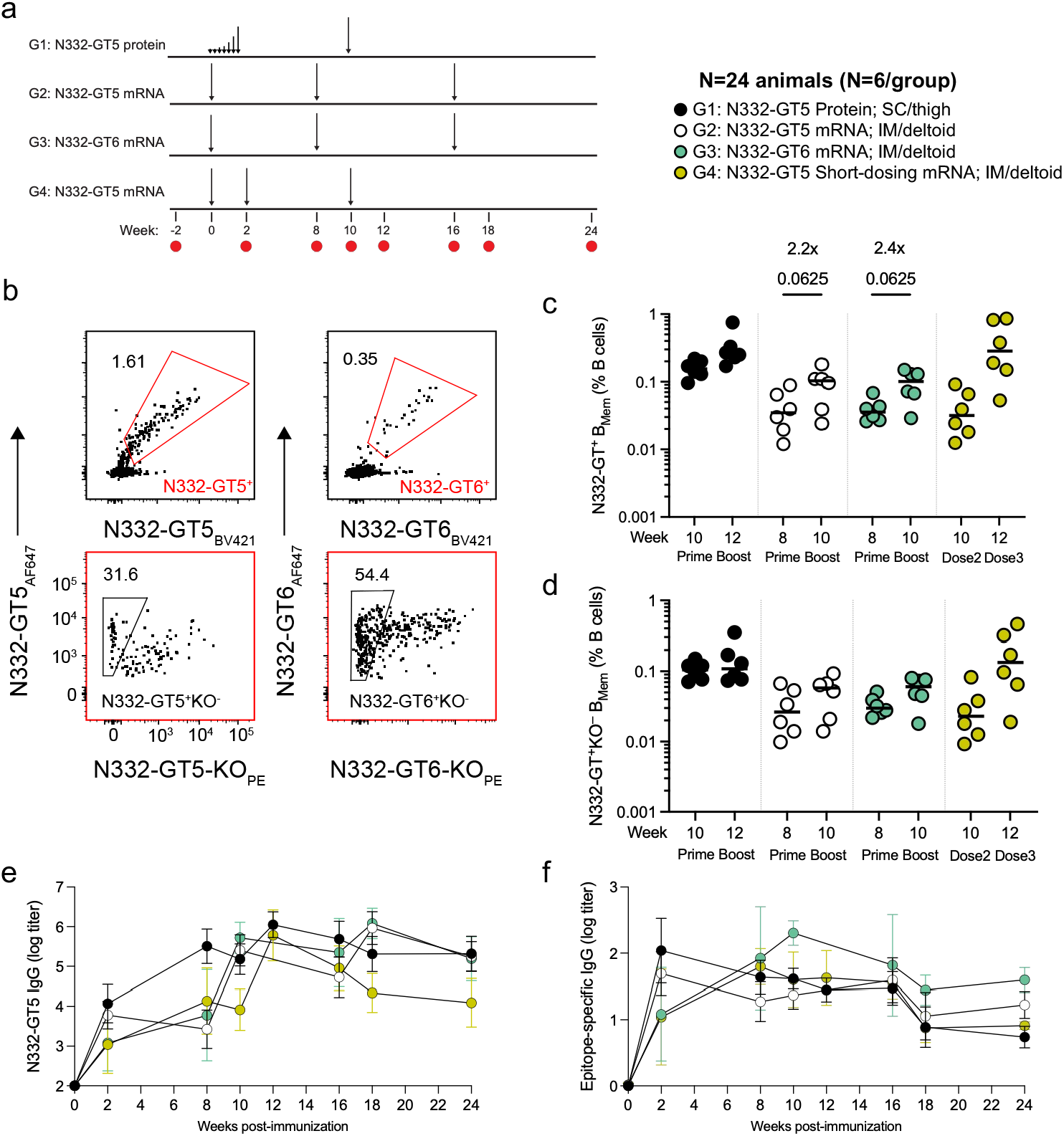
N332-GT6 mRNA primes and boosts epitope-specific B cell responses. **a,** Immunization schedule, in which animals (n=6/group) were immunized with N332-GT mRNAs or N332-GT5 protein at the indicated timepoints. Red dots indicate time-point of PBMC collection for B_Mem_ analysis. mRNA constructs are membrane-bound gp151. **b**, Flow cytometry gating of antigen-specific (N332-GT-AF647^+^N332-GT-BV421^+^) upper plots) and epitope-specific (N332-GT-AF647^+^N332-GT-BV421^+^N332-GT-KO-PE^−^) B_Mem_ cells (lower plots). Left, N332-GT5 immunized animal flow cytometry gating. Right, N332-GT6 immunized animal flow cytometry gating. **c**, Frequency of N332-GT^+^ B_Mem_ cells as a percentage of B cells. **d**, Frequency of N332-GT^+^KO^−^ B_Mem_ cells as a percentage of B cells. **e**, Time course of the endpoint titer of N332-GT5 binding IgG, as determined by ELISA. Geometric mean and SD are shown. **f**, Time course of the endpoint titer of N332-GT5 epitope-specific binding IgG, as determined by ELISA. Geometric mean and SD are shown.

N332-GT-binding B_Mem_ cells were analyzed post-prime and post-boost. N332-GT5 probes were used for analysis of Groups 1, 2 and 4, and N332-GT6 probes were used for analysis of Group 3 (**Supplementary Fig. 7**). N332-GT-binding B_Mem_ cells were detected after priming for all groups (**Fig. 2b-c**). N332-GT5 and N332-GT6 mRNA induced similar frequencies of N332-GT-binding B_Mem_ cells after the prime and after a homologous boost, and for both mRNAs the boost induced a modest but statistically non-significant increase in such B_Mem_ cells (**Fig. 2c**). The short-dosing regimen of N332-GT5 mRNA induced epitope-specific (N332-GT^+^KO^−^) B cell responses at frequencies comparable to that of N332-GT5 protein ED regimen after three doses of mRNA (**Fig. 2b, d)**. All groups induced anti-N332-GT5 and epitope-specific serum IgG antibody titers (**Fig. 2e-f, Supplementary Fig. 8**) with N332-GT6 inducing a stronger epitope-specific response than N332-GT5 post-boost (**Supplementary Fig. 8b**). In sum, these data demonstrated the ability of N332-GT mRNA immunogens to induce N332-GT-specific B_Mem_ cells in NHPs.

BG18-class precursor B cells are exceptionally rare in NHPs^24^. An estimated ∼50 total BG18-class naïve precursor B cell clones are predicted per adult animal^33^. To determine whether N332-GT5 or N332-GT6 mRNA elicited BG18-class cells, we sorted and sequenced N332-GT-binding B_Mem_ cells post-prime and post-boost. The previously established NHP definition of BG18 type I sequences was applied, defined by use of the rhesus macaque D gene *IGHD3-41* (homologous to human *IGHD3-3*) in the appropriate reading frame and presence of a GXXXXXE motif within the HCDR3, and an HCDR3 length of 20 amino acids or greater^24,28^. After priming, BG18 type I sequences were detectable in all animals (6/6) immunized with N332-GT6 mRNA (**Fig. 3a**). In contrast, only 1 of 6 N332-GT5 mRNA-immunized animals (Group 2) had detectable BG18 type I responses post-prime (**Fig. 3a**). N332-GT6 mRNA induced BG18-class B_Mem_ frequencies post-prime that were comparable to those from N332-GT5 protein ED immunization (Group 3 vs Group 1) and higher than from N332-GT5 mRNA (Group 3 vs. Group 2; *P* = 0.0087) (**Fig. 3b**).

**Fig. 3:**
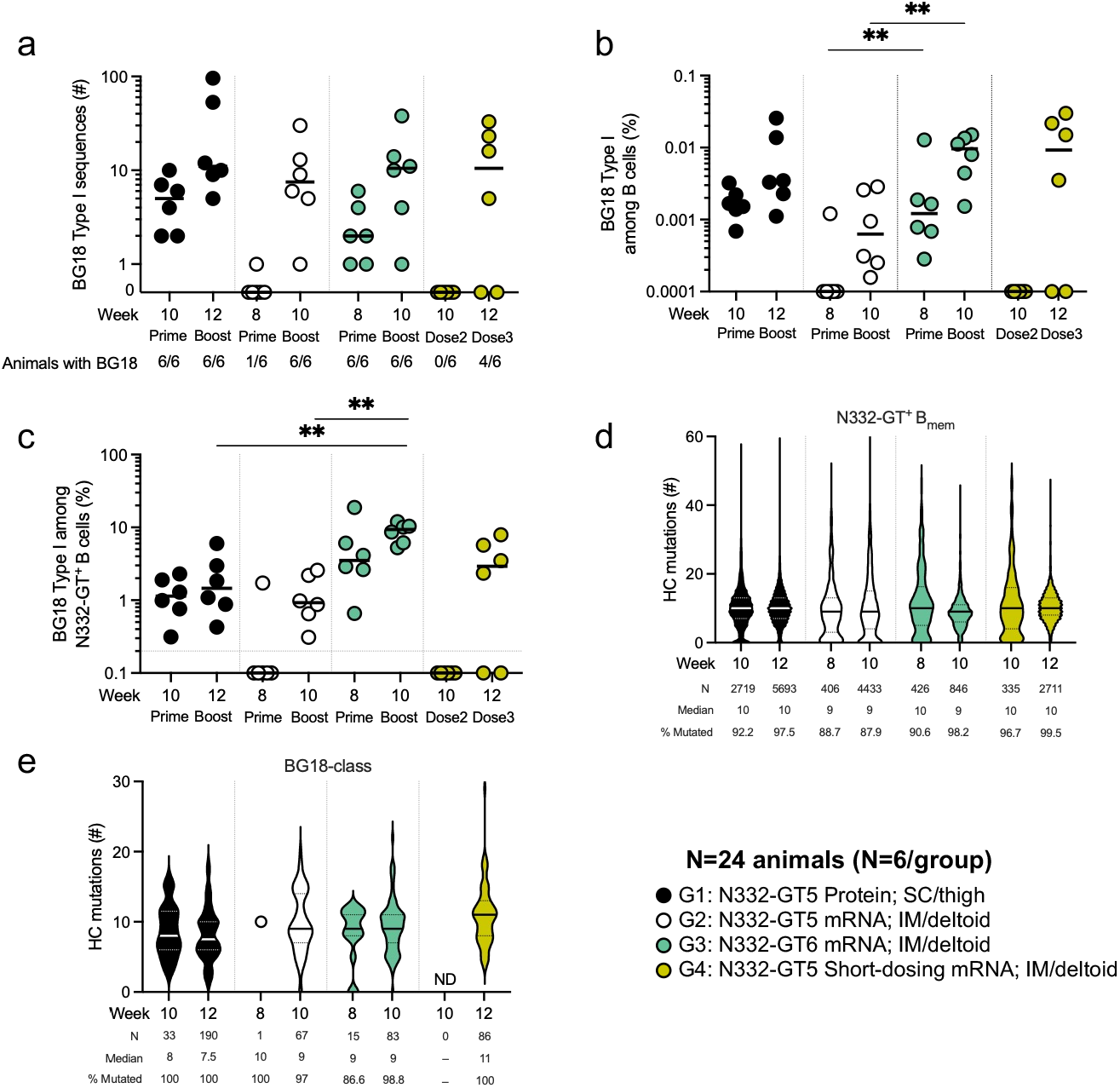
N332-GT6 mRNA is superior to N332-GT5 in priming and boosting BG18-class B cell responses. **a**, Number of BG18 type I sequences identified after priming and boosting. **b**, Percentage of BG18 type I sequences among B cells **c**, Percentage of BG18 type I sequences among N332-GT^+^ B cells. **d**, Number of heavy chain (HC) nucleotide mutations among N332-GT^+^ B cell sequences post-prime and post-booster immunization. **e**, Number of HC mutations among BG18 type I sequences post-prime and post-booster immunization. a-e, mRNA constructs are membrane-bound gp151.

After boosting, BG18-class responses were detected in all animals (6/6) in both Groups 2 and 3 (**Fig. 3a**). N332-GT6 mRNA induced substantially higher BG18-class B_Mem_ frequencies (15.3-fold) than N332-GT5 mRNA (Group 3 vs. Group 2, Week 10; *P* = 0.0087) (**Fig. 3b**). This was also true for BG18-class precursor B cells quantified as a percentage of antigen-specific cells, with a 10-fold enrichment in the N332-GT6 mRNA group (Group 3 vs Group 2; *P* = 0.0022) (**Fig. 3c**).

The short dosing regimen with N332-GT5 mRNA induced variable BG18-class B_Mem_ responses; 4 of 6 animals had detectable BG18 type I sequences (**Fig. 3b-c**).

Heavy chain SHM analysis of all N332-GT-binding B_Mem_ cells showed comparable numbers of mutations across groups (**Fig. 3d**). Almost all (∼87-100%) BG18-class B cells contained VDJ SHM post-boost, consistent with GC participation by BG18-class B cells (**Fig. 3e**). N332-GT5 and N332-GT6 mRNA immunization induced comparable numbers of HC mutations despite their ∼9-fold difference in affinity for BG18 type I precursors (**Fig. 3e**). Together, these data indicate that N332-GT5 and N332-GT6 mRNAs can efficiently prime BG18-class precursor B cells in NHPs, with N332-GT6 mRNA inducing more frequent detectable responses after a single immunization.

### Trimer stabilization, VLP display, and SC delivery do not improve N332-GT5 mRNA priming in NHPs

In parallel with NHP study 1, additional N332-GT5 mRNA constructs and delivery strategies were evaluated in NHPs to assess the effects of an Env-stabilizing mutation, an alternative antigen display format, and a different route of administration on immune responses. Four groups of animals were established. Group 1 received N332-GT5 mRNA-LNP administered IM and served as a control. Group 2 received N332-GT5 mRNA containing a stabilizing disulfide bond mutation (501C-663C; N332-GT5 1CC) to test whether additional trimer stability could limit trimer dissociation in vivo and thereby improve responses^52^. Group 3 received N332-GT5 mRNA co-delivered with HIV-1 *gag* mRNA to generate virus-like particles (VLPs) displaying multiple copies of membrane-bound N332-GT5, to test whether such antigen presentation could improve immunogenicity^53^. Group 4 received subcutaneous (SC) rather than IM immunization with N332-GT5 mRNA-LNP to test whether the route of administration could impact priming (**Fig. 4a**), as has been seen with adjuvanted protein^39^.

**Fig. 4:**
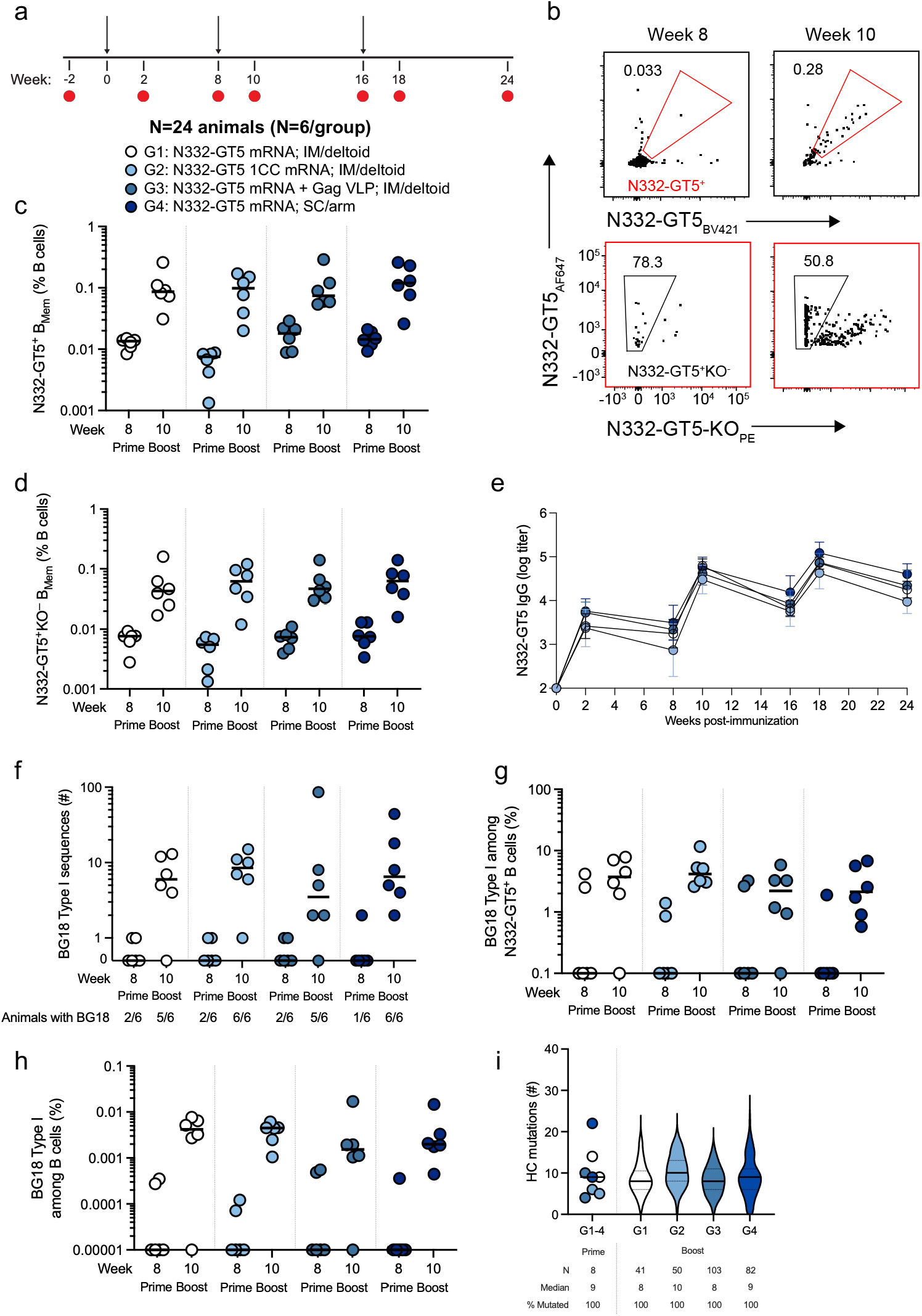
N332-GT5 mRNA constructs prime BG18-class B cells. **a**, Immunization schedule, in which animals (n=6/group) were immunized with different forms or routes of N332-GT5 mRNA at the indicated timepoints. mRNA constructs are membrane-bound gp151. **b**, Flow cytometry gating of antigen-specific (N332-GT5-AF647^+^N332-GT5-BV421^+^) memory B cells (B_Mem_) (upper plot) and epitope-specific (N332-GT5-AF647^+^N332-GT5-BV421^+^N332-GT5-KO-PE^−^) B_Mem_ cells (lower plot) post-prime (week8) and post-boost (week10). **c**, Frequency of N332-GT5^+^ B_Mem_ cells as a percentage of B cells. **d**, Frequency of N332-GT5^+^KO^−^ B_Mem_ cells as a percentage of B cells. **e**, Time course of the endpoint titer of N332-GT5 binding IgG, as determined by ELISA. Geometric mean and SD are shown. **f**, Number of BG18 type I sequences identified post-prime (week 8) and post-booster immunization (week 10) with N332-GT5 mRNA. **g**, Percentage of BG18 type I sequences among N332-GT5^+^ B cells. **h**, Percentage of BG18 type I sequences among B cells. **i**, Number of HC mutations among BG18 type I sequences post-prime (week 8) shown collectively across all groups and post-boost (week 10) shown for each individual group.

N332-GT5-binding B_Mem_ cells were quantified after the prime (week 8) and after the boost (week 10). N332-GT5-binding B_Mem_ cells were induced at similar frequencies across all four groups following both the prime and boost immunizations (**Fig. 4b-c**). Likewise, the frequencies of epitope-specific (N332-GT5^+^KO^−^) B_Mem_ cells were comparable among the groups (**Fig. 4b-d**). Consistent with these findings, N332-GT5-specific and epitope-specific serum IgG titers were similar across all four groups (**Fig. 4e, Supplementary Fig. 9**). BG18-class B_Mem_ cells were detected in 1-2 of 6 animals per group post-prime (**Fig. 4f**), consistent with N332-GT5 mRNA priming in NHP study 1 (shown in Figure 3) and emphasizing the reproducibility of the low rate of detectable BG18 precursor activation with N332-GT5 mRNA after a single prime. After homologous boosting with N332-GT5 mRNA, at least 5 of 6 animals in each group had detectable BG18-class B_Mem_, resulting in a total of 22 out of 24 (∼92%) animals with detectable BG18 type I responses after mRNA-LNP immunization (**Fig. 4f-h**). The frequencies of BG18-class B_Mem_ were similar across the four groups (**Fig. 4h**). All BG18-class sequences (100%) were mutated (**Fig. 4i**), indicating GC participation and selection under all four immunization conditions. Collectively, these data indicate that standard IM delivery of N332-GT5 mRNA effectively primed BG18-class precursor B cells without detectable improvement from additional trimer stabilization, VLP display, or SC administration.

### Affinities and specificities of BG18-class antibodies after priming and boosting with N332-GT mRNA in NHPs

Binding affinities were measured for monoclonal antibodies (mAbs) derived from BG18-type I B cell sequences isolated post-prime (week 8) and post-boost (week 10) from N332-GT5 and N332-GT6 mRNA immunized animals across both NHP studies (**Fig. 5**). To probe for differences in affinity maturation induced by N332-GT6 compared to N332-GT5, BG18-class mAbs derived from groups that had received N332-GT5-based immunizations were pooled and compared to mAbs derived from N332-GT6 immunized animals (**Fig. 3a**, **Fig. 4f**). BG18-class mAbs elicited by N332-GT5 or N332-GT6 exhibited high-affinity (sub-nM) binding to both N332-GT5 and N332-GT6 priming immunogens. BG18-class mAb binding was also assessed against a panel of increasingly native-like boosting candidates (B46, B50, B54, B11, and B20)^21,24,28^. B46, B50, and B54 are boost 1 immunogens that recently demonstrated success in a bnAb-inducing regimen in NHPs. B11 is a more native-like boost 1 candidate with only 6 GT mutations^21^, and B20 is a near-native trimer based on SF162p3 but with 3 GT mutations that was previously used as a boost 2 immunogen as part of a bnAb-inducing multi-boost regimen in NHPs^28^. Most post-prime (week 8) mAbs retained detectable binding across increasingly native-like boosters, although the percentage of mAbs with detectable affinity trended lower for the more native-like immunogens. Post-prime binding profiles were comparable between N332-GT5- and N332-GT6-induced mAbs. Post-boost (week 10) binding affinities showed similar binding affinities to N332-GT5, N332-GT6, and the booster candidates (**Fig. 5**). In sum, these data demonstrate that BG18-class antibodies elicited by a single vaccination with either N332-GT5 or N332-GT6 mRNA acquire high affinities for the priming immunogens and can recognize multiple downstream boosting immunogens, supporting the feasibility of sequential heterologous GT immunization strategies.

**Fig. 5:**
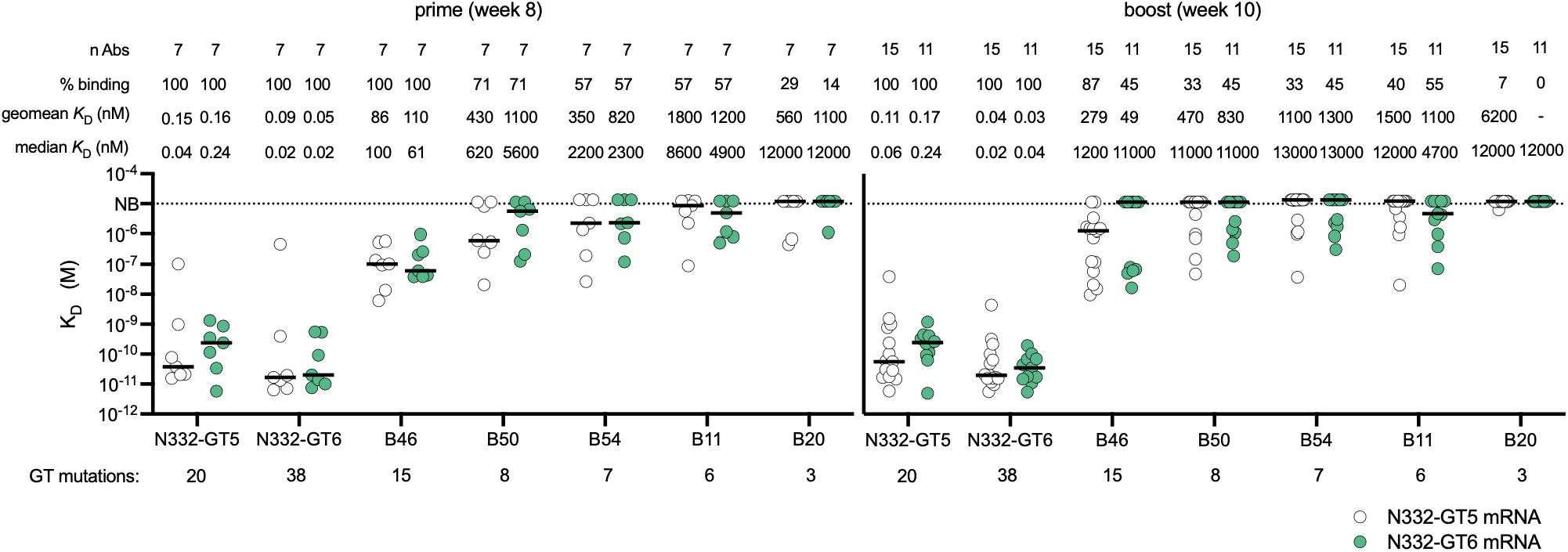
N332-GT5 and N332-GT6 induce BG18 type I precursors with similar affinities for boost candidates. Binding affinities (*K*_D_) of BG18 type I monoclonal antibodies post-priming immunization with N332-GT5 or N332-GT6 at week 8 and week 10 to N332-GT5/6 and different boosting candidates. Dashed line indicates the limit of detection. Bars represent the medians.

### Sequential heterologous boosting generates B cells recognizing increasingly native-like Env immunogens

In a third NHP study, two groups of animals were primed with N332-GT6 protein or mRNA and then given sequential heterologous booster immunizations to test for shepherding of BG18-class precursors towards wildtype HIV Env recognition (**Fig. 6a**). In the protein group, animals received ED priming with N332-GT6, followed by bolus immunizations with boost 1 immunogen B11 and boost 2 immunogen B20. All protein immunogens were soluble gp140 adjuvanted with SMNP. In the mRNA group, animals received a bolus immunization of N332-GT6 mRNA, followed by bolus immunizations with boost 1 immunogen B11 mRNA and boost 2 immunogen B20 mRNA. All mRNA immunogens were membrane-bound gp151.

**Fig. 6:**
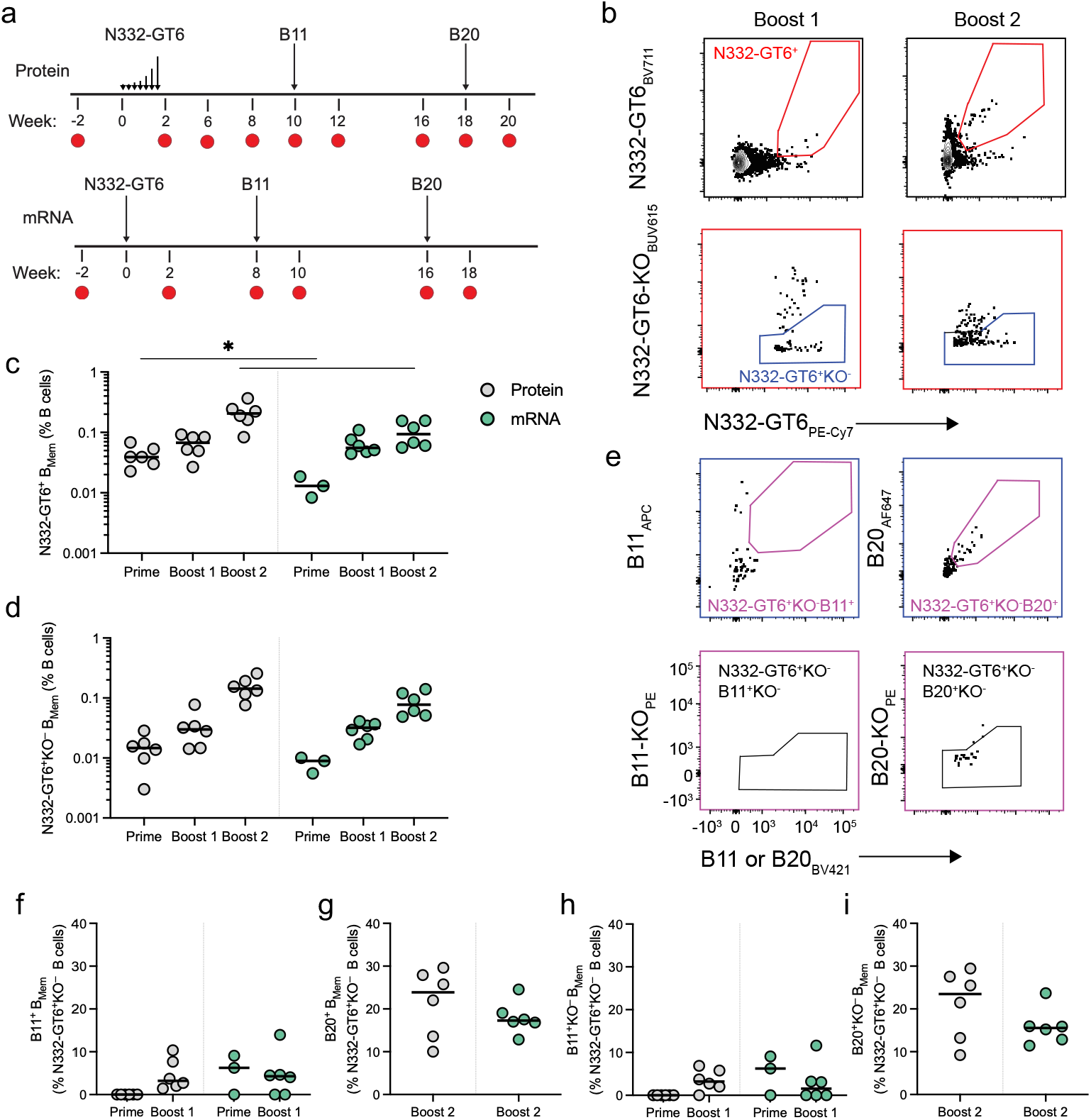
N332-GT6 mRNA priming induces development of cross-reactive booster antigen-specific B cells. **a**, Immunization schedule, in which animals (n=6/group) were primed with either N332-GT6 protein or mRNA, followed by heterologous booster immunizations of B11 and B20 at the indicated timepoints. mRNA constructs are membrane-bound gp151. **b**, Flow cytometry gating of antigen-specific (N332-GT6-PE-Cy7^+^N332-GT6-BV711^+^) B_Mem_ cells and epitope-specific (N332-GT6-PE-Cy7^+^N332-GT6-BV711^+^N332-GT6-KO-BUV615^−^) B_Mem_ cells. Left, post-B11 immunization. Right, post-B20 immunization. **c**, Frequency of N332-GT6^+^ B_Mem_ cells as a percentage of B cells. **d**, Frequency of N332-GT6^+^KO^−^ B_Mem_ cells as a percentage of B cells. **e**, Flow cytometry gating of cross-reactive antigen-specific (B11-APC^+^B11-BV421^+^ or B20-AF647^+^B20-BV421^+^) and epitope-specific (B11-APC^+^B11-BV421^+^B11-KO-PE^−^ or B20-AF647^+^ B20-BV421^+^ B20-KO-PE^−^) B_mem._ Left, post-B11 immunization. Right, post-B20 immunization. **f**, Frequency of B11^+^ B_Mem_ cells as a percentage of N332-GT6^+^KO^−^ B cells. **g**, Frequency of B20^+^ B_Mem_ cells as a percentage of N332-GT6^+^KO^−^ B cells. **h**, Frequency of B11^+^KO^−^ B_Mem_ cells as a percentage of N332-GT6^+^KO^−^ B cells. **i**, Frequency of B20^+^KO^−^ B_Mem_ cells as a percentage of N332-GT6^+^KO^−^ B cells. Lines represent median values. Statistical significance between groups was tested using unpaired two-tailed Mann-Whitney tests. *P ≤ 0.05; **P < 0.01; ***P < 0.001; ****P ≤ 0.0001. In c, d, f and h, for week 8 timepoint in the mRNA-immunized animals, PBMC samples from only three of six animals were available for flow cytometry analysis.

N332-GT6-binding B_Mem_ cells were quantified after each heterologous boost immunization (**Fig. 6b**). Higher N332-GT6-binding B_Mem_ cell frequencies were detected in protein-immunized animals compared to mRNA-immunized animals post-prime (*P* = 0.024) and post-B20 boost (*P* = 0.015), but similar frequencies were observed post-B11 boost (**Fig. 6c**). However, epitope-specific (N332-GT6^+^KO^−^) B_Mem_ cell frequencies were similar across all post-prime and post-boost timepoints between both groups, consistent with membrane-bound trimers reducing responses directed to the Env base^45,46^ (**Fig. 6d**). B_Mem_ cells binding the booster immunogens were also quantified (B11^+^ or B20^+^, **Fig. 6e**). Similar frequencies of B11 and B20 boost-binders were seen between protein and mRNA after B11 and B20 boosts, respectively (**Fig. 6f-i**). N332-epitope binding serum IgG titers using N332-GT6, B11 and B20 were measured (**Supplementary Fig. 10**). After N332-GT6 prime, epitope-specific IgG titers were higher in protein ED-immunized animals compared to mRNA bolus-immunized animals (protein ED week 10 vs. mRNA bolus week 8; *P* = 0.004) (**Supplementary Fig. 10b**). After B11 booster immunizations, B11-epitope binding IgG titers were detected at low levels in both groups, whereas B20 epitope-binding IgG titers were not detectable above background after B20 immunization (**Supplementary Fig. 10d, f)**. No neutralization of the wildtype heterologous viruses tested was detected at this time point, as expected. Serum from all animals post-B20 boost did neutralize a variant BG505 pseudovirus containing the N332-GT5 V1 loop (BG505_V1_GT5_)^24,27,28^ (**Supplementary Fig. 10g**). Overall, sequential heterologous immunization generated B cell responses capable of recognizing progressively more native-like Env booster immunogens.

### Sequential immunization with mRNA-LNPs guides BG18-class precursors to wildtype Env trimer binding

Epitope-specific (N332-GT6^+^KO^−^) B_Mem_ cells were sorted and sequenced after each heterologous prime-boost timepoint to identify BG18-class cells in the B_Mem_ cell pool. After the last heterologous booster B20, BG18 type I sequences were detected in 5 of 6 (83%) protein-immunized animals and 6 of 6 (100%) mRNA-immunized animals (**Fig. 7a**). BG18 type I sequences were detected for both groups at post-boost timepoints, for which a substantial number of epitope-specific B_Mem_ sequences were collected (e.g. at 2 weeks post-boost 1, 3027 BCR sequences were obtained from 12 animals across both groups). BG18 type I sequences were not detected post-prime, likely due to low sample cell viability and sequencing depth post-prime (54 BCR sequences from protein-immunized and 5 from mRNA-immunized animals, with only 2 mRNA-immunized animals available) (**Fig. 7a-c**). After Boost 1 (B11) and Boost 2 (B20) timepoints, BG18 type I B_Mem_ cell frequencies were similar between protein and mRNA groups (**Fig. 7b-c**). Thus, mRNA-LNPs were capable of boosting BG18-class precursor B cells in a heterologous boost sequential vaccination strategy.

**Fig. 7:**
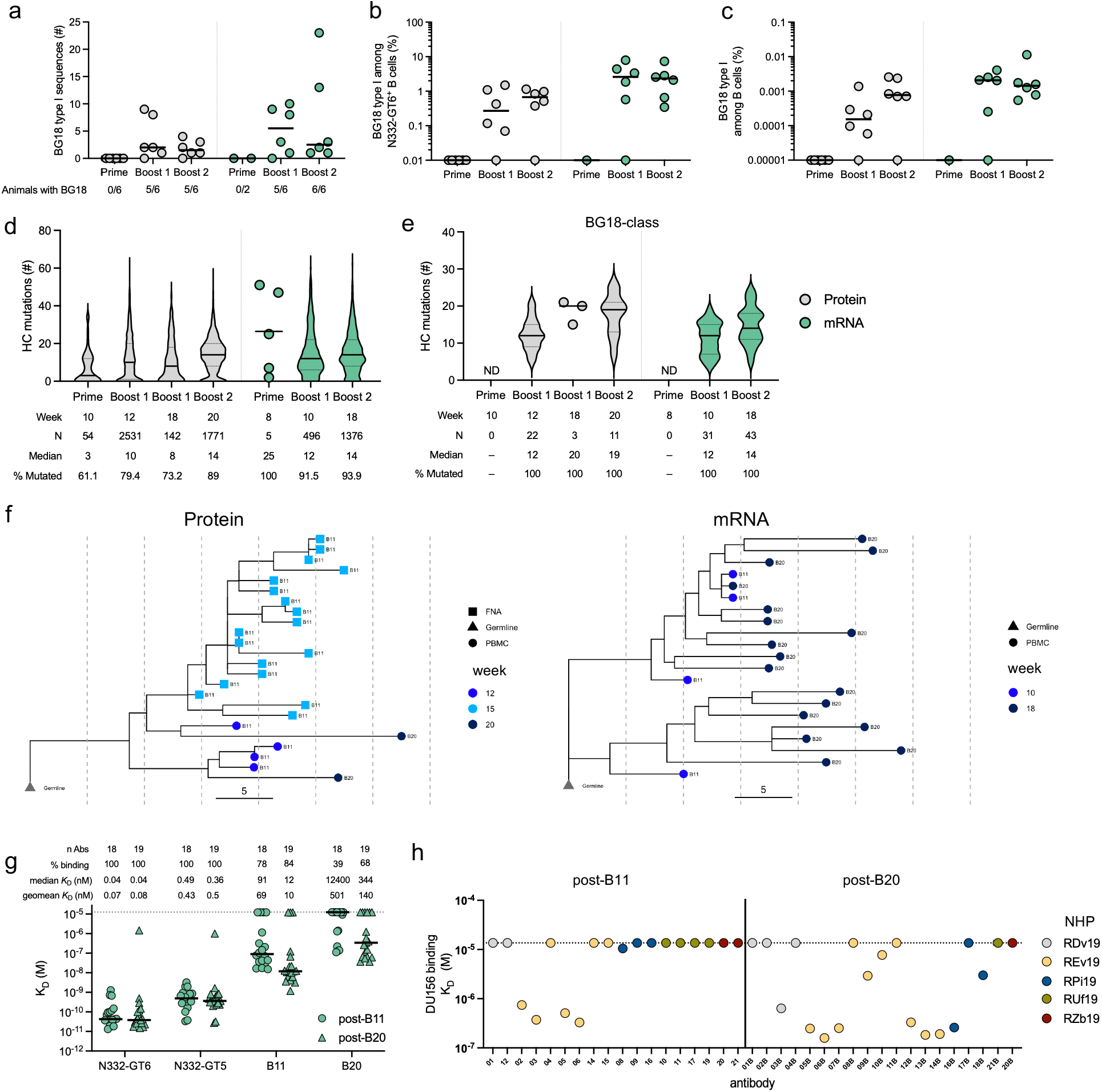
Heterologous prime-boosting by mRNA induces BG18-class B cells. **a,** Number of BG18 type I sequences identified post-N332-GT6 prime and post-B11 and post-B20 booster immunizations. **b,** Percentage of BG18 type I sequences among N332-GT6^+^ B cells. **c,** Percentage of BG18 type I sequences among B cells. **d,** Number of HC nt mutations among N332-GT6+ B cell sequences post-prime and post-booster immunizations. Data shown as individual points when fewer than 10 sequences were detected and otherwise as violin plots. **e,** Number of HC nt mutations among BG18 type I sequences post-prime and post-booster immunizations. Data shown as individual points when fewer than 10 sequences were detected and otherwise as violin plots. **f,** Clonal lineage trees of BG18 type I sequences recovered following booster immunizations. Top, clonal family of protein-immunized animal RDn19 in both the B_GC_ and B_Mem_ compartments at weeks 12 and 15 (from fine needle aspirates (FNA)) and week 20 (from PBMCs). Bottom, clonal family of mRNA-immunized animal REv19 in the B_Mem_ compartment at weeks 10 and 18. **g,** Binding affinities 2-weeks post-B11 and post-B20 booster immunizations to N332-GT5, N332-GT6, and booster immunogens B11 and B20 **h,** Binding affinities 2-weeks post-B11 and post-B20 booster immunization to wild-type trimers DU156. Dotted line across the plot represents the limit of detection.

Mutation analysis of the heavy chain in N332-GT6-binding antibodies revealed SHM in both groups (**Fig. 7d**). SHM among BG18 type I sequences was comparable after the first booster immunization in both mRNA and protein immunized animals (**Fig. 7e**). Every recovered BG18 type I sequence in both groups exhibited SHM. To understand maturation of heterologous boosted BG18-class B cells, we investigated clonal relationships of BG18-class precursor B cells in peripheral blood (both groups) and draining lymph nodes (protein only). Clonal lineage analysis of both protein and mRNA groups suggested continuing maturation of primed BG18-class B cells following heterologous boosters (B11 and B20) (**Fig. 7f**). This indicates N332-GT6 priming can activate ultra-rare BG18-class precursors and those BG18-class B cells are capable of being further affinity matured following heterologous boosters.

To test whether N332-GT6 was required to prime BG18-class B cell precursors or could be bypassed in favor of a more native booster, animals were primed with B11 in a separate small NHP study (Study 4). Priming with B11 alone did not induce detectable BG18-class B cell precursors, in contrast to priming with N332-GT6 (**Supplementary Fig. 11**).

Several BG18-class clonal lineages were primed and expanded following the first mRNA heterologous boost in NHP Study 3 (week 10; median of 3 clones. **Supplementary Fig. 12a**). The number of clones primed by a single N332-GT6 mRNA immunization is difficult to accurately determine because BG18-class clones were rare, raising the possibility of undersampling. Nevertheless, in NHP study 1, most animals had more than one clone recovered after N332-GT6 mRNA priming (week 8; median of 2 clones. **Supplementary Fig. 12b**). The number of BG18-class clonal lineages detected after homologous boosting with N332-GT5 or N332-GT6 mRNA was 3-4 across both NHP studies 1 & 2 (**Supplementary Fig. 12b-c**).

BG18 type I sequences isolated from the mRNA sequential immunization group were expressed as mAbs and tested for binding affinities to N332-GT5, N332-GT6, B11, and B20 (**Fig. 7g)**. BG18-class mAbs displayed high-affinity binding to both priming and heterologous Env immunogens (**Fig. 7g)**. The mAbs were then tested for binding to wildtype Env trimers. Notably, after a single heterologous boost with B11 mRNA, BG18-class BCRs from one expanded lineage bound the wild-type Env trimer DU156, and additional clones acquired wildtype trimer binding after the B20 boost (**Fig. 7f, h, Supplementary Fig. 13**). Overall, sequential mRNA-LNP immunization successfully expanded multiple BG18-class B cell lineages and enabled maturation of a subset toward wildtype Env trimer recognition.

### Structure of an mRNA-induced BG18 type I antibody that binds wildtype Env trimers

We identified a BG18 type I lineage with low (<5%) SHM that had acquired affinity for a heterologous wildtype trimer following mRNA vaccination with only one GT prime and one heterologous boost (N332-GT6 and B11 mRNA) (**Supplementary Fig. 14**). Cryo-electron microscopy (cryo-EM) was applied to characterize the binding mode of a mAb from this lineage isolated after the B20 boost (REv19_06B). In this complex (which included the CD4 binding site-specific VRC01-class bnAb 3BNC117 for orientation) REv19_06B Fab targets the V3-glycan epitope as anticipated (**Fig. 8a, Supplementary Fig. 3**). The REv19_06B HCDR3 shows high structural similarity to the human BG18 bnAb (**Fig. 8b-c**), consistent with other BG18 type I antibodies^4,24,28^. GT mutations in B20 make contacts to the tip of the HCDR3 of REv19_06B Fab (**Fig. 8c**). Unlike previously described BG18 type I antibodies, the variable domain is twisted relative to BG18, which rotates the LC away from the V1 loop (**Fig. 8d-e**), in contrast to the classical BG18 binding mode in which the LC straddles the V1 loop (**Fig. 8d**). The orientation of the REv19_06B LC away from the V1 loop may contribute to this lineage achieving wildtype binding with less affinity maturation, as the V1 loop glycans and sequence diversity typically present an obstacle for the LC. However, a key step in BG18 bnAb maturation is the acquisition of contacts with the N332 glycan, which this antibody had not yet achieved after the B20 boost.

**Fig. 8:**
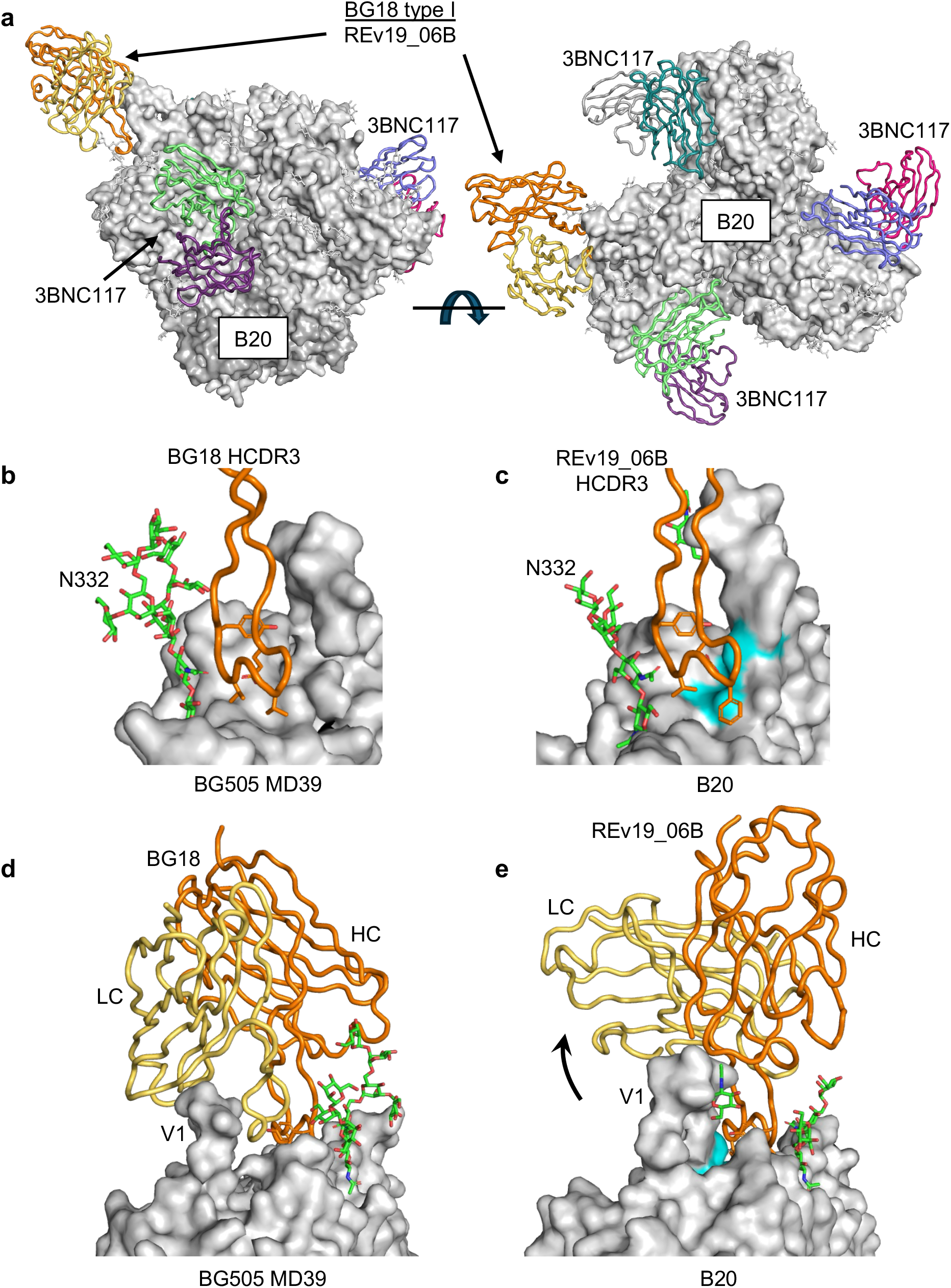
Minimally mutated BG18-class antibody structural interactions with near-native trimer B20. **a**, Structure of Env trimer B20 in complex with elicited NHP BG18 type I Fab. The B20 trimer is shown in gray surface representation, BG18 type I Fab Rev19_06B is shown in orange HC and yellow LC. The complex includes three Fabs of the CD4 binding site bnAb 3BNC117. **b,** BG18 in complex with native-like Env trimer BG505 MD39 (PDB ID 6DFG) with close up view of the HCDR3. **c**, Rev19_06B Fab in complex with B20 with close view of the HCDR3. The 3 GT mutations in B20 are shown as cyan surface color. **d**, Structure of BG18 in complex with BG505 MD39 showing the BG18 binding mode with the LC straddling the V1 loop. **e**, The Rev19_06B binding mode showing the LC rotated away from the V1 loop.

## Discussion

GT vaccines can selectively activate ultrarare naïve bnAb-class B cells and guide their maturation despite unfavorable competitive dynamics in the naïve B cell repertoire. A recent study demonstrated that GT sequential vaccination can induce potent serum and B_Mem_ bnAb activity targeting the Env V3-glycan site in NHPs, using adjuvanted protein immunogens and escalating-dose priming^28^. Other studies have demonstrated the promise of sequential immunization strategies in NHPs for the V2-apex epitope, which has more favorable B cell precursor frequencies and affinity maturation pathways in NHPs than in humans^7,54^. mRNA-LNP technology can accelerate clinical development of immunogens, facilitate multi-component vaccines, and support delivery of natively membrane-anchored and glycosylated multimeric antigens, all of which are favorable attributes for HIV vaccine development^20,21,45,46,55,56^. Key open questions were whether the induction of bnAbs by GT vaccination in NHPs was unique to the vaccination modalities implemented, and whether mRNA-LNP technology is suitable for priming ultra-rare B-cell precursors and eliciting V3-glycan type bnAb-class B cell responses. Here we show, across three separate NHP studies, two critical findings: mRNA-LNP versions of both N332-GT5 and N332-GT6 consistently induce BG18-class bnAb-precursor B cells; and a three-step, 4-month, mRNA-LNP sequential vaccination regimen can generate wildtype Env-binding bnAb-class B_Mem_.

A key design goal was to develop an immunogen with higher affinity for “true” BG18 precursors and then evaluate priming of those ultrarare precursors in the NHP model system. During the design process we also selected mutations that substantially improved the expression of soluble N332-GT6. With both affinity and expression enhanced, N332-GT6 mRNA showed superior BG18-class B_Mem_ responses after just a single priming immunization (6/6 animals primed), prior to any booster immunization. Improved binding of N332-GT6 to BG18-class precursors may facilitate better selection dynamics and stronger signaling to drive clonal expansion, allowing for detection of BG18-class B_Mem_ often prior to a booster immunization. Together, across all three NHP studies reported here, N332-GT5 and N332-GT6 mRNA successfully primed BG18-class B cells in ∼92% (44 of 48) of animals after homologous or heterologous mRNA immunizations, which is a remarkable achievement for such rare B cells.

Sequential mRNA-LNP immunization drove affinity maturation of BG18-class B cells to recognize near-native (B20) and native (DU156) Env trimers after only 2 or 3 immunizations, highlighting the potential of the mRNA-LNP platform for GT sequential vaccination strategies. Future studies should define an optimal affinity window for productive boosting. The range of binding affinities of the primed precursor B cells to heterologous boosters may play a critical role in shepherding, where excessively high or low affinities might be detrimental^23,24,57^. Evidence that antibody feedback can affect homologous booster outcomes in mice was recently reported^58^. Potential antibody feedback limitations in NHPs were observed after N332-GT5 immunization, wherein high titers of epitope-specific serum IgG following an escalating dose prime were proposed to have led to faster antigen clearance and epitope masking of the BG18 epitope, resulting in suboptimal boosting responses with homologous antigen^33^. Here, evaluating boosting with two sequential heterologous immunogens, (B11 and B20), we demonstrate that by maintaining an antigenic distance between each sequential immunogen, an mRNA-LNP encoded sequential vaccination GT regimen can successfully shepherd responses towards BG18-class bnAb development, potentially reducing the negative impact of antibody feedback. Overall, rational design of GT immunogens and boosting regimens considering factors like Env backbone, antigen affinity, and precursor frequency will be key to improving shepherding outcomes.

These studies had some limitations. We elected to test shepherding using three total immunizations, with a N332-GT6 prime and B11 and B20 boosts. More than half of the post-B20 BG18-class B_Mem_ Abs expressed bound wildtype Env trimer DU156; however, DU156 neutralization was not detected. Human clinical trial data for GT shepherding of VRC01-class bnAbs suggest an affinity threshold of 50-100 nM K_D_ is needed for neutralization activity^49^. The binding affinities elicited in this study were close to, but did not exceed, the 100 nM K_D_ affinity threshold. Additional heterologous boosters may be needed to determine whether the observed BG18-class lineages could mature to neutralize wild-type viruses. Lastly, it remains untested whether two or more homologous priming immunizations are beneficial compared to a single homologous prime in a full sequential vaccination regimen.

In sum, this study demonstrates key advances for GT vaccine strategies using the mRNA-LNP platform in the context of HIV vaccine development, and provides a framework for designing complex, heterologous immunization regimens that can guide rare B cell precursors toward bnAb maturation.

## Methods

### Adjuvants

Saponin QS-21 was purchased from Desert King and used as received. Phosphorylated HexaAcyl Disaccharide (PHAD^®^) MPLA was obtained from Avanti. GMP-process QS-21 SMNP was prepared using a tangential flow filtration-based detergent dilution process, as previously described^59^. Briefly, SMNP components were solubilized in 20% (w/v) MEGA-10 surfactant at a 10:2:1:1 mass ratio of QS-21 saponin:cholesterol:MPLA:dipalmitoylphosphatidyl choline at batch size of 500 mg of QS-21. This starting detergent/lipids solution was diluted with phosphate buffered saline (PBS, pH 7.4) to reach a final QS-21 concentration of 5 mg/mL and MEGA-10 concentration of 7.5%. The adjuvant/lipid mix mixture was then stepwise diluted with PBS with sufficient incubation time between each dilution step to allow for controlled SMNP self-assembly. After SMNP assembly, the sample was concentrated 20-fold on a 100 kDa hollow fiber TFF membrane (mPES, surface area: 0.16 m2, Repligen). After concentration, the sample underwent continuous diafiltration for 10 diafiltration volume equivalents against PBS pH 6.5. The sample was then filtered through a 0.2 μm membrane and used to fill 3 mL 2R Type I USP borosilicate glass vials with 0.6 mL of sample. Vials were closed with 13 mm bromobutyl 4023/50, Flurotec coated, B2-40 grey stoppers and 13 mm aluminum caps with green plastic flip-off. Quil-A SMNP was produced as previously described^37^. Protein-immunized animals received QS-21 SMNP or Quil-A SMNP adjuvant. QS-21 SMNP has shown comparable immune responses to Quil-A SMNP in NHPs when immunized with Env trimer MD39 protein^37^.

### Protein production

Biotinylated trimers, His-tagged trimers and untagged trimers were expressed and purified from 293F or Expi293 cells (Thermo Fisher Scientific) as described previously^28^. Untagged N332-GT5 was produced as described previously^60^.

### Design of N332-GT6

Lentivirus-based mammalian display directed evolution was performed as described previously^4^. A random mutagenesis library was first screened that covered the majority of the BG18 epitope (residues 114 to 366 in HxB2 numbering) including the V1 and V3 loops. This library was screened against antibodies pre14, preD, preF, preG, preI and VL3-21.

A structure-guided combinatorial library that sampled the following AA was assembled: 139R/T, 141I/M, 150I/M, 153Q/E/L/V, 155K/M, 178K/R, 302N/T, 303T/S, 323M/L, 326T/M/A/V, 327K/R/M, 328M/V, 337T/S. This library identified mutations E153Q and T303S. This library was screened against antibodies VL3-19, preD, preI, pre12, pre14.

A second structure-guided library was assembled encoding for the following AA: 389(all 20AA), 412H/Q/D/E, 442(all 20AA), 444N/K/S/R. The library was screened against antibodies preD and pre14.

Next, two saturation mutagenesis libraries were assembled targeting segments spanning residues 120 to 212 and 290 to 446. These libraries were screened against antibodies pre12, pre14, preD, preI, LV3-21 as well as antibody pairs pre20+preD, pre27+pre14, pre36+preI, VL3-21+pre14.

The most enriched mutations from these libraries were incorporated into two combinatorial libraries. Combo 1: 124Q/P, 125H/L/Y/F, 127N/T/I/D/A/V, 130H/Q, 159L/F, 166R/W, 179R/L, 185fR/W, 186N/I/H/L, 188N/K/D/E, 195N/I/H/L.

Combo 2: 307I/M/L, 319D/A/Y/S, 320M/I/V/L/F, 346A/V, 367A/G, 371R/M/G/V, 372T/S, 424I/L/V, 428H/Q/P, 430S/I. These combo libraries were screened against antibodies pre20+preD, pre27+pre14, VL3-21+pre14, which identified 11 more mutations that were incorporated into 29mutA to make N332-GT6.

### Antibody production

Antibodies were produced at Genscript as described previously^28^.

### Cell surface antigenic profile

HEK293F cells were transfected with different DNA-encoded, membrane-bound trimers using 293Fectin in Opti-MEM. DNA (25 µg) and 293Fectin (50 µL) were diluted separately in 833 µL Opti-MEM and incubated for 5 min at room temperature. The DNA solution was then added to the 293Fectin solution and incubated for an additional 15 min at room temperature. The resulting transfection complexes were added to HEK293F cells (25 mL at 1 × 10⁶ cells/mL) and incubated for 2 days. For flow cytometry, 1.8 mL of transfected cells were collected by centrifugation at 650 × g for 5 min at 20°C and stained with the corresponding primary Fabs (100 µL at 10 µg/mL in FACS buffer consisting of HBSS, 1 mM EDTA, and 1% BSA) for 1 h at 4°C. Cells were washed twice and then stained with SYTOX Green (1:1,000) and Alexa Fluor 647 anti-human IgG F(ab′)₂ (1:5,000) for 20 min at room temperature. Following two additional washes, cells were resuspended in 150 µL FACS buffer and analyzed by flow cytometry using a NovoCyte 3000 with NovoSampler Pro (Agilent ACEA). Approximately 50,000 live cells were collected per well. FlowJo 10 was used to identify single, viable cells based on forward and side scatter (FSC/SSC), doublet exclusion, and SYTOX Green fluorescence. Alexa Fluor 647 (APC-A) fluorescence was quantified as the median fluorescence intensity (MFI) and analyzed using GraphPad Prism. Fab binding to each membrane-bound trimer was compared across constructs to characterize and compare their antigenic profiles.

### Thermal stability

Differential scanning calorimetry experiments were performed on a MicroCal VP-Capillary differential scanning calorimeter (Malvern Instruments) as previously described^23^.

### Antigenic profile

Biolayer interferometry was used to characterize the antigenic profile of the immunogens as previously described^28^.

### HCDR3 junction frequency

BG18 class precursors were identified by searching a NGS data set of 1.1×10^9^ human BCR heavy chains^4,50^ for sequences with the same length HCDR3 (23 aa), D3-3 in the same reading frame and position and JH6 usage. Among the identified BG18 precursors, the frequency of all aa at the D3-3+2 position was determined.

### mRNA formulation

mRNA-LNPs were synthesized in vitro and passed quality control tests as previously described (Steward-Jones et al STM 2023). The secretion signal sequence from the pHLsec vector (MGILPSPGMPALLSLVSLLSVLLMGCVAETG)^61^ was used for all constructs. For Gag immunizations, we used mRNA encoding SynGagIIIB^62^ with a mutation (V370A) introduced that was located in the p2 linker region and was selected for in patients with protease inhibitor (bevirimat) resistant strains^63,64^.

### Animals and immunizations

#### Mouse models and subject details

Healthy adult male or female mouse models homozygous for the Pypre5 and Pypre7 were generated and maintained in the C57BL/6J (CD45.2^+/+^) background. Pypre5 and Pypre7 express authentic BG18 germline heavy chains identified by NGS sequencing, accompanying with the same BG18 inferred germline (iGL) light chain (PMID: 31672916). 8–14-week-old male B6.SJL-Ptprca Pepcb/BoyJ mice (CD45.1^+/+^) from The Jackson Laboratory (Bar Harbor, ME) were purchased and used in this study. Mice had free access to food and water, controlled temperature and a 12:12 h light-dark cycle and were housed at the animal facility of Mass General Hospital (MGH). The mouse maintenance and experiments were performed in accordance with the approved protocols by the Institutional Animal Care and Use Committee (IACUC) of MGH, an Association for Assessment and Accreditation of Laboratory Animal Care (AAALAC) International-accredited facility, under Animal Study Protocols 2016N000286 and 2016N000022.

#### B cell adoptive transfer in mice

This assay was carried out as previously reported^21^. Lymphocytes from Pypre5 and Pypre7 mice were isolated from spleens. Freshly collected spleens were crushed in FACS buffer (DPBS containing 2% FBS) and filtered with a 70 μm cell strainer, after which the cell suspension was subjected to pan B isolation kit (Miltenyi Biotec) according to the manufacturer’s protocol. Finally, the B cell pellets were suspended in DPBS. Cell viability and concentration were analyzed by the cell counter (Luna), then naive B cells were transferred to CD45.1^+^ recipient mice at the specified frequencies via intravenous (IV) injection. The Pypre5 and Pypre7 cells were transferred at the cell number of 5K (low), 50K (intermediate) and 200K (high), respectively.

#### Mouse Immunization

This assay was carried out as previously reported^21^. Protein immunogens were diluted in PBS at the volume of 100 μL/per mouse and were mixed with Sigma adjuvant dissolved in PBS at the volume of 100 μL/per mouse. Then the immunogen/adjuvant complex was shaken on the shaking table for 30 min before each mouse was administrated with 200 μL immunogen/adjuvant complex via intraperitoneal (IP) injection. mRNA lipid nanoparticle (LNP) immunogens were diluted in PBS at the volume of 100 μL/per mouse. Each mouse was given 100 μL mRNA immunogen via intramuscular (IM) injection in the thigh muscles of the two side hind limbs. The immunogen dosage used is noted in the schematic graphs in each figure.

#### Flow cytometry of murine cells

This assay was carried out as previously reported^21^. Single cell suspensions were generated by mechanical dissociation of spleen (post protein immunization) or inguinal, popliteal and iliac lymph nodes (post LNP-mRNA immunization). Red blood cells were removed by ACK lysing buffer and then cell pallets were suspended in fluorescence-activated cell sorting (FACS) buffer (DPBS containing 2% FBS). Cell viability was stained with Live/Dead Blue in PBS at 4°C for 10 min and then cells were Fc blocked (clone 2.4G2, BD Biosciences, 1:100 dilution) in FACS buffer at 4°C for another 10 min. Cells were preincubated with freshly fluorescently labeled protein probes for 20 min before the cocktail of antibodies against surface markers merged and stained for another 20 min at 4°C. Afterward, samples were washed once and filtered before flow cytometry recording. Samples were acquired and events were recorded on a BD LSR Fortessa for flow cytometry analysis. Data was analyzed by FlowJo software (Waters Biosciences).

#### Cell sorting of murine cells

This assay was carried out as previously reported^21^. Spleens were collected from naïve Pypre5 and Pypre7 mice and crushed and filtered, then cell pallets were suspended in FACS buffer. The samples were proceeded with Pan B cell isolation kit II in accordance with manufacture’s protocol. After that, cells were incubated with an antibody cocktail against surface markers (Dump (Ly6G/Ly-6C-CD3-F4/80)^-^B220^+^/IgM^+^ or IgD^+^) and anti-mouse CD45 hashtag oligos for 30 min at 4°C. Afterwards, cells were thoroughly washed and suspended in FACS buffer with DAPI. Cell sorting was performed on a BD FACS Aria Fusion at a flow rate of <5000 events/second for tube sorting using a 70 μm nozzle. Recorded events were analyzed by FlowJo software and sorted samples were used for the 10x Genomics single cell experiment below for naïve B cell repertoire characterization.

#### BCR sequencing and analysis of murine cells

This assay was carried out as previously reported^21^. ∼5000 targeted live cells were sorted from each sample and no more than 12,000 in total per reaction were encapsulated and converted into several DNA libraries following the 10X Next GEM Single cell 5’ protocol (10X Genomics). Briefly, single cells were isolated with Gel Bead-In-EMulsions (GEMs) using the Chromium controller provided by 10X genomics, resulting in uniquely barcoded transcriptome for each individual cell. After initial cDNA amplification and conversion to dsDNA, individual sequencing libraries were generated for gene expression, V(D)J repertoires and hashtag oligos. Library quality was analyzed using a Tapestation 4200 (Agilent). Libraries were pooled at a ratio based on depth requirements established by 10X genomics and subsequently sequenced using a Nextseq 2000 sequencer (Illumina). Raw base call files generated by sequencing were demultiplexed, aligned and aggregated using the pipeline offered as part of Cell Ranger (10X Genomics). For V(D)J repertoire analysis, immunoglobulin v genes were determined by Cell Ranger followed by a customized analysis pipeline that includes the Pypre5 and Pypre7 heavy chain and BG18 iGL light chain sequences.

#### Rhesus macaque studies

For the N332-GT5 versus N332-GT6 mRNA study (NHP study 1), rhesus macaques (*Macaca mulatta*) were housed at Alpha Genesis Primate Research Center and maintained in accordance with NIH guidelines. The study was approved by the Alpha Genesis IACUC. All animals were between 2.8 to 3.3 years of age and between 3.2 and 5.1 kg at the time of the first immunization. Each immunization group (n = 6/group) contained at least two male animals. All doses at their respective timepoint were given bilaterally, split between the left and right side. Group 1 animals were primed SC with a total of 100 µg soluble N332-GT5 plus 750 µg of Quil-A SMNP in the thighs, following a seven-dose 12-day escalating dose strategy^40^. The dose for each of the 7 injections per side of N332-GT5 were: D0: 0.10 µg, D2: 0.22 µg, D4: 0.58 µg, D6: 1.58 µg, D8: 4.28 µg, D10: 11.65 µg, D12: 31.60 µg, totaling 50 µg per side and 100 µg after bilateral administration. The dose for each of the 7 injections of Quil-A SMNP per side were: D0: 0.75 µg, D2: 1.61 µg, D4: 4.35 µg, D6: 11.81 µg, D8: 32.10 µg, D10: 87.38 µg, D12: 237.00 µg, totaling 375 µg per side and 750 µg after bilateral administration. Animals were then boosted with a bolus immunization of 100 µg soluble N332-GT5 plus 750 µg SMNP at week 10. Groups 2, 3, and 4 all received bolus mRNA immunizations, which were administered IM in the deltoids. Group 2 animals were immunized with 100 µg N332-GT5 gp151 mRNA, Group 3 received 100 µg N332-GT6 gp151 mRNA, and Group 4 received 100 µg N332-GT5 gp151 mRNA. For Groups 2 and 3, animals were immunized at weeks 0, 8 and 16. For Group 4, animals received a shorter dosing regimen where immunizations were performed at weeks 0, 2, and 10.

For the N332-GT5 mRNA study (NHP study 2), Indian rhesus macaques were housed at the Emory National Biomedical Research Center (ENBRC) and maintained in accordance with NIH guidelines. The study was approved by the Emory University Institutional Animal Care and Use Committee (IACUC) under the protocol 202200104. All animals were between the ages of 2.5 to 3.8 years and between 3.5 to 7.8 kg at the start of the first immunization. RMs were pair housed for the duration of the study. Each immunization group (n = 6/group) contained at least one female animal. RMs were immunized bilaterally according to their respective study group with the dose split between the left and right deltoids, as follows: (i) Group 1 with a total dose of 100 µg N332-GT5 gp151 mRNA administered intramuscularly (IM), (ii) Group 2 with a total dose of 100 µg N332-GT5 1CC gp151 mRNA given IM, (iii) Group 3 with 100 µg N332-GT5 gp151 plus 50 µg *gag* mRNA given IM, and (iv) Group 4 with 100 µg N332-GT5 gp151 mRNA given SC. All animals were immunized at three timepoints at weeks 0, 8 and 16.

For the heterologous prime-boost N332-GT6 study (NHP study 3), RMs were housed at the ENBRC and maintained in accordance with NIH guidelines. The study was approved by the Emory University IACUC under the protocol 202100053. All animals were between 3 to 3.5 years of age, between 4.2 to 5.7 kg at the time of the first immunization and consisted of all males. For the protein group, six NHPs were given immunizations SC split between the left and right mid-thighs. At priming, NHPs were immunized with a total of 100 µg soluble N332-GT6 plus 375 µg QS-21 SMNP using a seven-dose 12-day escalating dose strategy^40^ at week 0. The dose for each of the 7 injections per side of N332-GT6 were: D0: 0.10 µg, D2: 0.22 µg, D4: 0.58 µg, D6: 1.58 µg, D8: 4.28 µg, D10: 11.65 µg, D12: 31.60 µg, totaling 50 µg per side and 100 µg after bilateral administration. The dose for each of the 7 injections of QS-21 SMNP per side were: D0: 0.375 µg, D2: 0.805 µg, D4: 2.175 µg, D6: 5.905 µg, D8: 16.05 µg, D10: 43.69 µg, D12: 118.5 µg, totaling 187.5 µg per side and 375 µg after bilateral administration. At boosting, NHPs received a bolus immunization totaling 100 µg BG505-B11 plus 375 µg QS-21 SMNP at week 10 and 100µg SF162p3-B20 plus 375 µg QS-21 SMNP at week 18. For the mRNA group, six NHPs were given immunizations IM split between the left and right deltoids. At priming, NHPs were immunized with a bolus immunization of 100 µg N332-GT6 gp151 at week 0. At boosting, NHPs received a bolus immunization of 100 µg BG505-B11 gp151 at week 8 and 100 µg SF162p3-B20 at week 16.

For the B11 mRNA-LNP priming study (NHP Study 4), rhesus macaques were housed at the Wisconsin National Primate Research Center. All procedures were approved by the Wisconsin National Primate Research Center Institutional IACUC and maintained in accordance with NIH guidelines. Six animals between 2-6 years of age with equal numbers of females and males were immunized with a single dose of BG505-B11 gp151 mRNA-LNP administered SC. Animals had previously been immunized with an unrelated GT immunogen (10E8-GT12) targeting 10E8-class MPER responses and had no detectable BG18-class B cell responses prior to the BG505-B11 immunization^27^.

### Lymph node fine needle aspiration in NHPs

Lymph node fine needle aspirations (LN FNAs) were performed on protein-immunized animals to sample the left and right draining LNs, which were identified by palpation and performed by a veterinarian. A 22-gauge needle attached to a 3-mL syringe was passed into the LN up to five times. Samples were transferred into RPMI media containing 10% FBS and 1x penicillin and streptomycin. Ammonium-chloride-potassium (ACK) lysing buffer was used if the sample was contaminated with red blood cells. LN FNA samples were frozen down in freezing media composed of 10% DMSO in FBS and stored in liquid nitrogen until analysis.

### Flow cytometry and sorting of NHP cells

Frozen FNA samples or peripheral blood mononuclear cells (PBMCs) were thawed and recovered in RPMI media containing 10% FBS, supplemented with 1x penicillin/streptomycin and 1x GlutaMAX. The recovered cells were counted and plated in a 96-well U-bottom plate up to 5 x 10^6^ cells per well, stained with 1:25 Fc block (BD Biosciences). Fluorescent antigen probes were prepared by mixing fluorophore-conjugated streptavidin (SA) with small amounts of biotinylated protein antigen probes (N332-GT5, N332-GT6, B46, BG505-B11, SF162p3-B20, “wild-type” (WT) probes, and their respective “knockout” (KO) probes) in a step-wise addition by adding one-third of the SA to the biotinylated protein at a time with a 15-minute incubation in between. Cells were first incubated with the KO probes for 20 minutes and then with N332-GT5 or N332-GT6 WT probes for 15 minutes in the dark at 4°C and stained with surface antibodies according to **Supplementary Tables 1-3**. Cells were then washed twice with FACS buffer (2% FBS in PBS). For samples being sorted, anti-human Total-Seq-C oligo-tagged anti-β2M and anti-CD298 hashtag antibodies (2 µL for 5 million cells; BioLegend) were added to each individual sample at the same time as the surface antibody mix to multiplex animal samples. All samples were acquired and sorted on a FACSymphony S6 (BD Biosciences).

For the heterologous prime-boost N332-GT6 study at week 8, PBMC samples from only three of six animals in the mRNA-immunized group were available for analysis.

### BCR sequencing and analysis of NHP cells

N332-GT^+^ or N332-GT^+^KO^−^ B_Mem_ cells were sorted into 1.5 mL tubes with RPMI media supplemented with 50% FBS. After sorting, cells were washed once and loaded onto the Chromium GEM-X chip and controller following the manufacturer’s instructions (10x Genomics). The indexed V(D)J, Feature Barcode and Gene Expression libraries of sorted cells were prepared following the protocol for Chromium GEX-X Single Cell 5’ Kit v3 (10x Genomics). Custom primers were designed to target rhesus macaque BCR constant regions (**Supplementary Table 4**). Forward primers were used at a final concentration of 1 µM and reverse primers at 0.5 µM, each per 100 µL of PCR reaction. All libraries were sequenced on a NovaSeq6000 (Illumina).

Cell Ranger v8.0.0 was used for generating FASTQ files, performing full-length V(D)J read assembly, and processing feature barcode and gene expression data. The V(D)J reads were assembled in de novo in Cell Ranger and then aligned to a custom *Macaca mulatta* germline VDJ reference using the Change-o package v1.3.0 within the Immcantation Portal^65^ v4.4.0. Sequences were de-multiplexed by the hashtag read counts using the MULTI-seqDemux command in Seurat^66^ v5. Inferred germline sequences were added with CreateGermline.py within the Change-o package. The germline D gene sequences were masked from analysis since these cannot be accurately predicted. For all analyses, paired HC-LC sequences were used. Somatic hypermutation analysis was performed using the observedMutations command within the SHazaM package^65^ v1.2.0. The number of mutations for each heavy chain was determined by counting the number of nucleotide changes between the observed sequence and the inferred germline sequence. Amino acid mutation frequencies were calculated using a custom Python script from in-frame translated BCR heavy chain sequences aligned to their corresponding translated germline references (germline_alignment_d_mask), excluding masked positions. Clonal lineages for each animal were identified using the DefineClones.py command from the SHaZam package, applying clustering thresholds determined by the disToNearest function. Sequences where VH genes aligned to alleles IGHV3-10001, IGHV3-10001_S4205, IGHV3-10001_S4375, IGHV3-3601_S5206, IGHV3-3601_S6650, IGHV3-NL_1101_S5714, IGHV4-79-a, IGHV4-NL_101_S0419 were found to have unusually high levels of nucleotide substitutions across all time points relative to their inferred germline sequences. This was likely due to incorrect V-gene assignment stemming from an incomplete V(D)J reference library. Therefore, these sequences were excluded from subsequent analyses. Clones with paired HC-LC BCR sequences were used when building clonal trees. Maximum-likelihood lineage trees were built for clonal families using Dowser^67^ with the pml method within the GetTrees function. For the lineage trees, the branch length indicates the estimated number of total HC mutations and its most recent common ancestor in lineage.

BG18 type I sequences were identified by a HCDR3 length of at least 20 amino acids long, use of the rhesus macaque D gene *IGHD3-41* in-frame, and the presence of a GXXXXXE motif, as previously described^28^.

### ELISA

Antigen-specific antibody binding was measured by ELISA using Corning™ 96-well half-area plates (Cat. #3690) with either anti-His (NHP study 2 & 3 experiments) or streptavidin (NHP study 1) antigen capture. For anti-His capture, plates were coated with anti-His antibody (2 µg/mL; GenScript) for 2 h at room temperature (RT), blocked with PBST (0.2% Tween-20) containing 5% skim milk and 1% FBS for 1 h, and incubated overnight at 4°C with His-tagged antigen (2 µg/mL). For streptavidin capture, plates were coated with streptavidin (2 µg/mL) for 2 h at RT, blocked overnight at 4°C with 2% BSA in PBS-T (0.2% Tween-20), and incubated with biotinylated antigen (0.5 µg/mL) for 1 h at RT.

Following antigen capture, plates were washed three times and incubated with serially diluted serum or monoclonal antibodies for 1 h at 37°C and 80% humidity. Plates were then incubated for 1 h at RT with HRP-conjugated anti-human IgG (Jackson ImmunoResearch, Cat. #709-035-149) at 1:5,000 dilution. After washing, TMB substrate (Thermo Fisher, Cat. #002023) was added for 6–8 min and the reaction was stopped with 0.5 M H₂SO₄. Absorbance was measured at 450 and 570 nm using a VersaMax plate reader, and background-corrected values were calculated by subtracting the 570-nm signal from the 450-nm signal. Endpoint titers were determined by interpolation from four parameter logistic curves using graphpad prism 11. The endpoint was determined by plotting OD values against the dilution factor and fitted by nonlinear regression using a four-parameter logistic model in Graphpad prism 11. The endpoint titer was defined as the reciprocal of the dilution at which the fitted curve crossed OD of 0.1 (∼3x background), obtained by interpolation.

### Surface plasmon resonance

For the SPR in **Fig 5**, **Fig. 7** and **Supplementary Fig. 12**, binding kinetics were measured on a Carterra LSA using HC30M or CMDP Sensor Chip as described previously^28^. Binding experiments in **Fig. 1b** were performed on a BIacore 8K instrument using Sensor Chip PEG Series S (Cytiva) and 1x HBS-EP+ pH 7.4 running buffer (20x stock from Teknova, Cat. H8022) supplemented with BSA at 1 mg/mL. We followed Biacore control software instructions to prepare chip surfaces for ligand capture. In a typical experiment, approximately 500 RU of capture antibody (His-tag Rabbit pAb, Gnescript Cat. No. A00174) in 10 mM Sodium Acetate pH 4.5 was amine coupled in both flow cells. Phosphoric acid 1.7% was our regeneration solution with 30 sec contact time and injected three times per cycle. Ligands were purified His-tagged HIV trimers. The capture contact time was 90 sec, and trimer concentration was 1 µg/mL in running buffer. The ligand capture levels were between 50 and 100 RU. Raw sensograms were analyzed using Biacore Insight Evaluation software (Cytiva), reference spot and blank double referencing, Langmuir model. Fab concentrations were quantified on NanoDrop 2000c Spectrophotometer using absorption signal at 280 nm. For the experiments in **Supplementary Fig. 1d** and **Supplementary Fig. 5b**, binding was measured using the ProeOn XPR36 (Bio-Rad) using the HC30M XanTec chips by capturing His-tagged trimer (ligand) with a His-tag antibody (pAb, rabbit, Genscript, catalog no. A00174) and flowing Fab (analyte)^24^.

### CryoEM sample preparation, imaging, and data processing

The following complexes were prepared in microcentrifuge tubes and left to incubate overnight at 22°C: 1) 0.2 mg SF162P3-B20 SOSIP + 0.2 mg Rev19_06B Fab + 0.2 mg 3BNC117 Fab; 2) 0.2 mg MD65 N332-GT6 SOSIP + 0.3 mg RM20A3 Fab. Each complex was purified on a GE AKTA Pure FPLC system equipped with a Cytiva HiLoad 16/600 Superdex 200 pg gel filtration column. Peak corresponding to complexes were pooled and concentrated using Millipore Amicon 100 kDa MWCO centrifugal filters. The final concentrations as determined by A280 on a Thermo Fisher Scientific NanoDrop with extinction coefficient (1% absorbance) 15.5 were 6.7 mg/mL (SF162P3-B20 complex) and 4.7 mg/mL (MD65 N332-GT6 complex). Lauryl maltose neopentyl glycol (LMNG; Anatrace) at a final concentration of 0.005 mM was mixed with the sample and 3 µL of the solution was immediately loaded onto UltrAuFoil R. 1.2/1.3 (Au, 300-mesh; Quantifoil Micro Tools GmbH) grids. Before sample application, grids were glow-discharged (Pelco easiGlow; Ted Pella) at 15 mA for 25 sec. Grids were prepared using a Vitrobot Mark IV (ThermoFisher Scientific) with the chamber set to 4°C with 100% humidity and variable blot times between 3–6 sec.

SF162P3-20 complex grids were loaded into a TFS Glacios microscope operating at 200 keV and equipped with a TFS Falcon 4 direct electron detector (190,000x magnification, resulting in 0.725 Å/pixel), and MD65 N332-GT6 complex grids into a TFS Glacios 2 microscope operating at 200 keV and equipped with a TFS Falcon 4i direct electron detector (190,000x magnification, resulting in 0.718 Å/pixel) (**Supplementary Table 5**).

EPU software (Thermo Fisher Scientific) was used for automated data collection. Micrograph movie frames were motion and CTF corrected using cryoSPARC Live^68^. The exposures were transferred to cryoSPARC where the remaining data processing was performed. Micrographs were filtered using GCTF parameters and particles were subsequently picked using the template picker and extracted. During extraction, particles were downscaled to 0.932 Å/pixel (SF162P3-B20 complex) or 1.034 Å/pixel (MD65 N332-GT6 complex) to reduce box size and increase the speed of downstream jobs. The extracted particles underwent three rounds of 2D classification to remove noisy particle picks. After 2D classification, a 3D ab initio reconstruction was performed to generate an initial volume for future refinement steps. Non-uniform refinement^69^ was performed with global CTF refinement. Specific imaging, collection, and data processing details for individual structures can be found in Supplementary Fig. 3 and Supplementary Table 5.

### CryoEM model building and refinement

Model building was performed by docking an initial model of N332-GT5 SOSIP in complex with RM20A3 Fab and a BG18-like Fab (PDB 8T49) into the N332-GT6 complex map using ChimeraX^70^ and mutating residues to match the correct sequence in Coot^71^ 0.9. RM20A3 Fv coordinates were retained from the initial model. Initial models of SF162P3-B20 SOSIP and Rev19_06B Fv were generated using AlphaFold 3^72^ and docked into the respective cryoEM map. Coordinates for 3BNC117 were taken from PDB 4JPV. Iterative rounds of manual model refinement in Coot and real space refinement in Phenix^73^ were used to refine models into the cryoEM maps. Final models were evaluated using MolProbity and EMRinger in the Phenix software suite. Refinement statistics are summarized in Supplementary Table 5. All maps and models have been deposited to the Electron Microscopy Data Bank (EMDB) and Protein Data Bank (PDB), respectively (see Supplementary Table 5 for accession codes).

### Site-specific glycan analysis

Using DeGlyPHER^74^, the glycoprotein was digested with Proteinase K, then sequentially deglycosylated with Endo H followed by PNGase F, and the resulting peptides with signature mass modifications were analyzed by LC-MS/MS as previously described^24^.

### Statistical analysis

Statistical tests were performed in GraphPad Prism 10. To assess the significance between groups in their respective studies, we conducted unpaired two-tailed Mann-Whitney tests. For comparison of *K*_D_s in Fig. 1b, and Supplementary Fig. 1d, we performed a paired *t* test on log-transformed *K*_D_ values. P values are defined throughout as follows: not significant, P > 0.05; *P ≤ 0.05; **P < 0.01; ***P < 0.001; ****P ≤ 0.0001.

## Data availability

All data supporting the conclusions of this study are presented in the main figures and supplementary materials. The BCR sequencing data generated in this study will be deposited in the NCBI Gene Expression Omnibus (GEO). The LC-MS/MS data from the glycosylation analysis will be deposited in the MassIVE repository. The 3D cryo-EM reconstructions have been deposited in the Electron Microscopy Data Bank (EMDB) under accession codes EMD-79016 and EMD-79015. The corresponding atomic models have been deposited in the Protein Data Bank (PDB) under accession codes 38OQ and 38OP.

## Acknowledgments

We thank Melissa Ferguson and the veterinary staff at Alpha Genesis Inc. for conducting NHP experiments. We thank Eva Rakasz and veterinary staff at the Wisconsin National Biomedical Research Center for conducting NHP experiments. We thank Diane Cheon for her assistance with mouse generation. We are also grateful to the Flow Cytometry and Next Generation Sequencing core facilities at the La Jolla Institute for Immunology for their services. This work was supported by National Institute of Allergy and Infectious Diseases (NIAID) UM1 AI100663 (Scripps Center for HIV/AIDS Vaccine Immunology and Immunogen Discovery, CHAVI-ID), UM1 AI144462 (Scripps Consortium for HIV/AIDS Vaccine Development, CHAVD) (to F.D.B., D.J.I., A.B.W., G.S., J.M.S., S.C., and W.R.S.), R61 AI183471 (to P.R.R.), U42 PDP11023 and P51 OD011132 to Emory National Biomedical Research Center, the Bill and Melinda Gates Foundation Collaboration for AIDS Vaccine Discovery (NAC INV-007522, INV-008813, and INV-034657 to W.R.S. and A.B.W.), IAVI support to W.R.S., and the IAVI Neutralizing Antibody Center (NAC) to W.R.S and A.B.W. The NovaSeq 6000 used in this work was funded by NIH grant S10OD025052.

## Author Contributions

Conceptualization: SC, WRS, PRR, JMS, FDB

Methodology: JMS, PRR, IP, BLF, ZX

Investigation: IP, BLF, AL, AS, ZX, SP, OK, PJM, EMZ, JRW, CAC, KMM, SB, CTF, CK, DL, NP, DG, YA, RT, EG, JT, AK, XH, VRL, JTN, XL, JKD, UN, TS, PL, DGC, GO, PRR, JMS

Visualization: SC, WRS, PRR, JMS, IP, BLF, AS

Funding acquisition: FDB, DRB, DJI, ABW, GS, PRR, JMS, SC, WRS

Project administration: SH, DGC, SC, WRS, PRR, JMS

Supervision: FDB, JCP, JRY, ABW, GS, SC, WRS, PRR, JMS

Writing – original draft: PRR, JMS, IP, BLF, SC, WRS

Writing – review & editing: All authors

