## Supplemental Figures for "HIV broadly neutralizing antibody precursor maturation induced by mRNA vaccination in nonhuman primates"

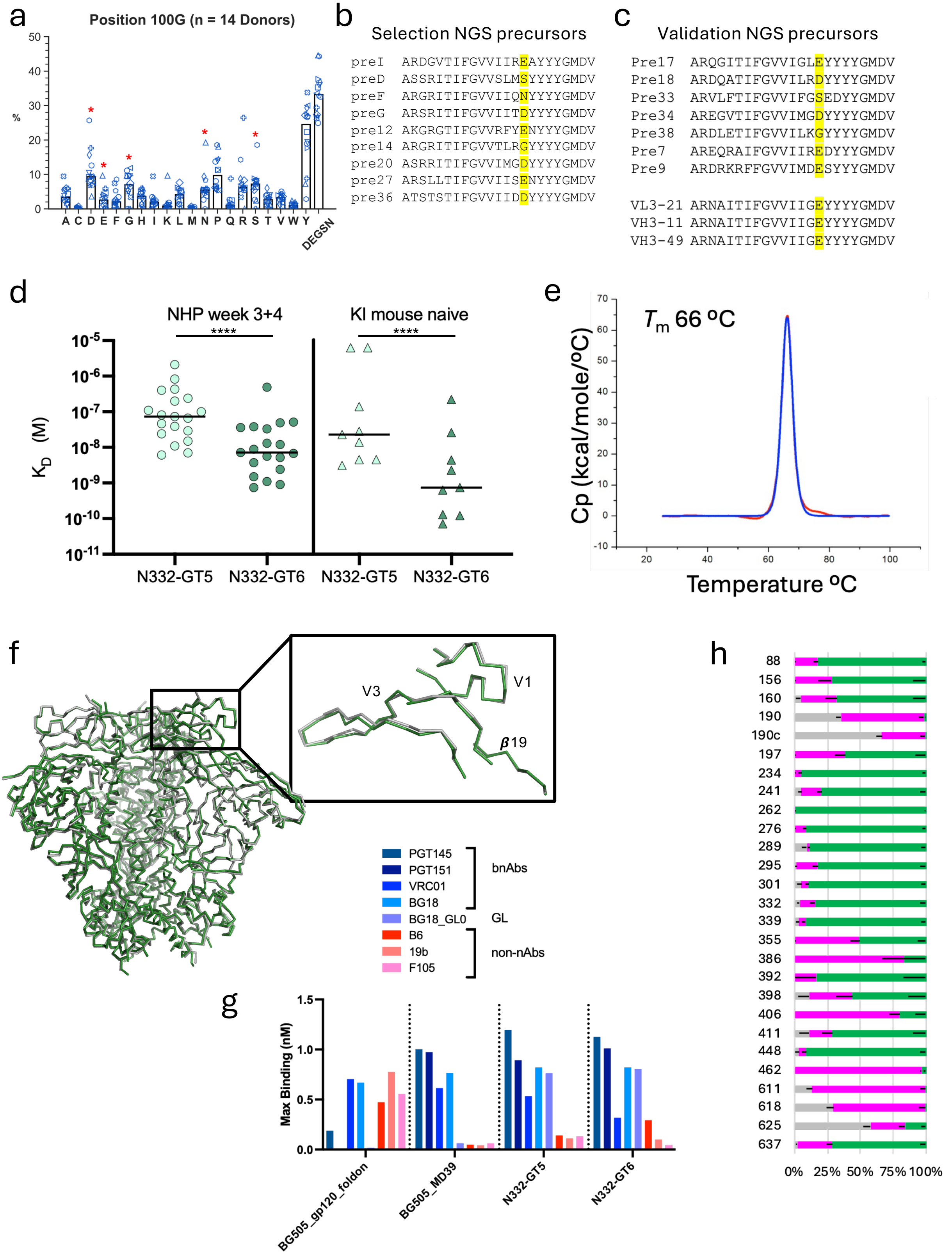


**Supplementary Fig. 1**

**Supplementary Fig. 1: N332-GT6 characterization**

**a,** Frequency of amino acids at the 100G position in BG18 type I precursors identified in an NGS dataset of antibody HCs. Asterisk indicates amino acids that have been shown to bind N332-GT immunogens with Glu being the most favorable.

**b,** HCDR3 sequence of BG18 type I precursors used for selection reagents in directed evolution experiments.

**c,** HCDR3 sequence of BG18 type I precursors used for immunogen validation.

**d,** Affinity of N332-GT5 and N332-GT6 for BG18 type I precursors isolated from NHPs 3 and 4 weeks post-N332-GT5 prime or from naïve mouse B cells in the BG18 iGL HC knock-in mouse.

**e,** Melting temperature of N332-GT6.

**f,** Structural alignment of unliganded N332-GT6 (green) and N332-GT5 (gray) with zoom view of the BG18 epitope.

**g**, Antigenic profile of N332-GT5 and N332-GT6 with positive control (BG505_MD39) and negative control (BG505_gp120_foldon) measured by BLI.

**h**, Glycan profile of N332-GT6. Green indicates high mannose glycans, pink indicates complex type glycans and gray indicates unoccupied PNGS.

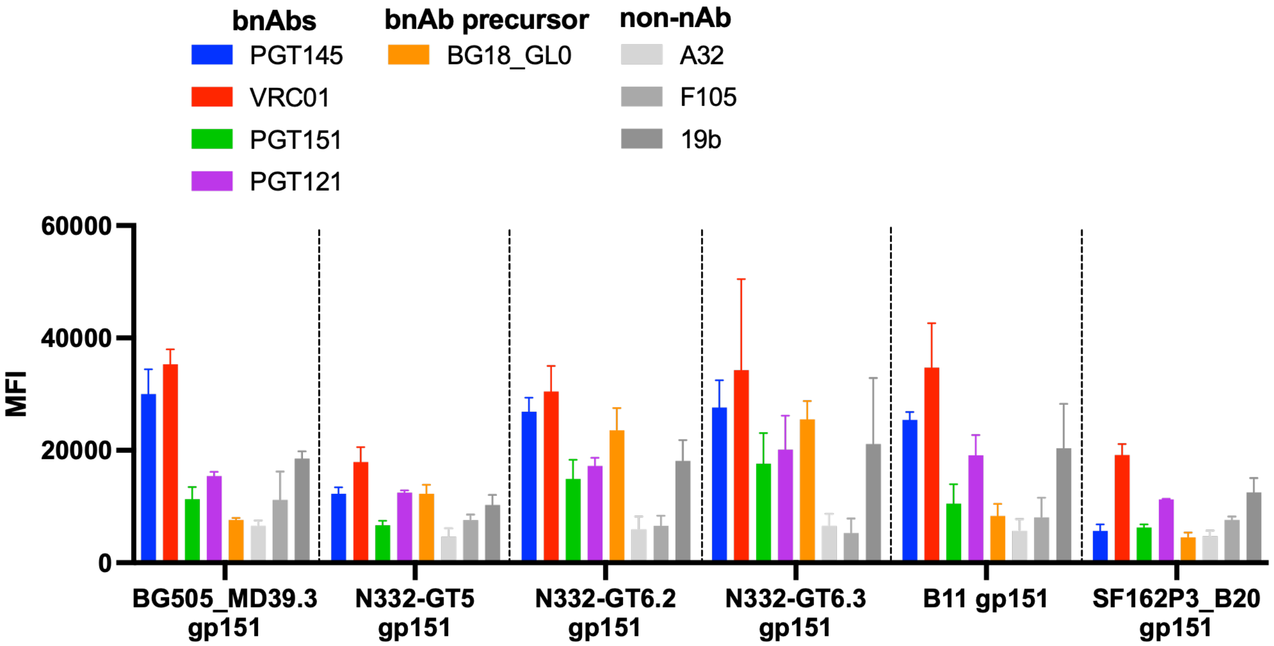


**Supplementary Fig. 2**

**Supplementary Fig. 2: Cell surface antigenic profile of immunogens**. The N332-GT immunogens were expressed on the cell surface by transient transfection of DNA in 293F cells and 2 days later the antigenic profile was measured by FACS analysis.

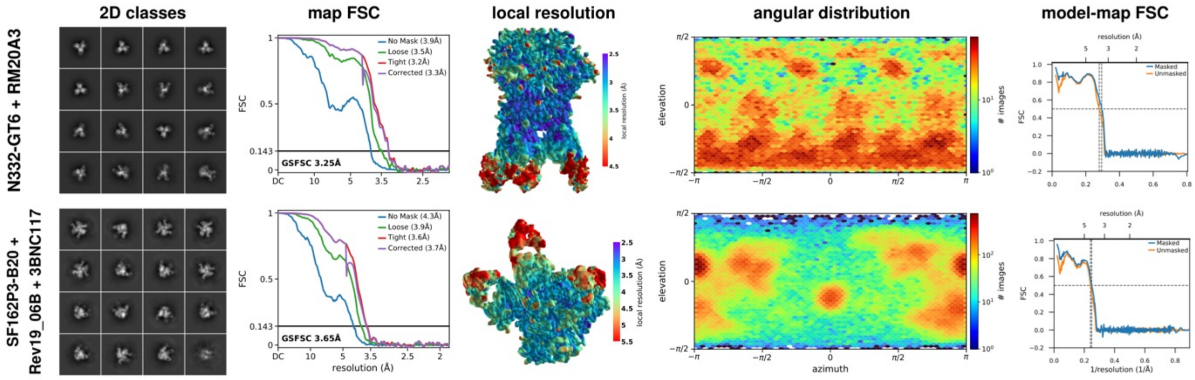


Supplementary Fig. 3

**Supplementary Figure 3: CryoEM data processing statistics.** Representative 2D class averages, gold standard Fourier shell correlation (FSC) resolution calculations (0.143 cutoff), map colored by local resolution (0.143 FSC), angular distribution of particle images, and atomic model to map FSC (0.5 cutoff).

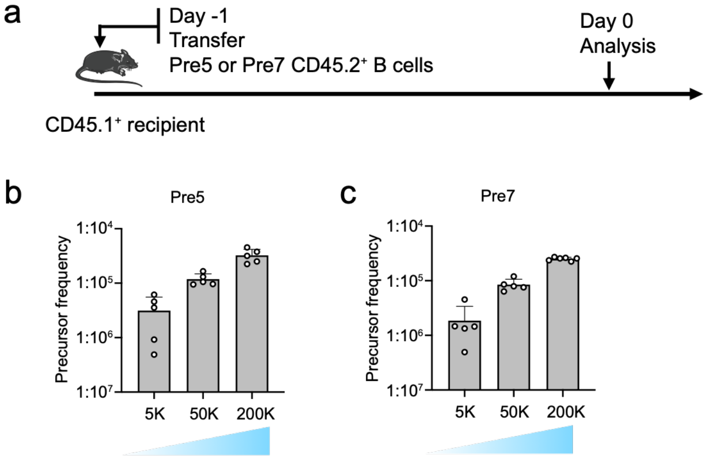


Pre5 HCDR3: ARDKDTIFGVVIIGEYYYYGMDV

Pre7 AREQRAIFGVVIIREDYYYGMDV

**Supplementary Fig. 4**

**Supplementary Fig. 4: Pre5 and Pre7 precursor frequency counting**

**a**, Schematic showing naïve CD45.2 Pre5 and Pre7 B cells at the number of 5K, 50K or 200K adoptively transferred into CD45.1 WT mice (day -1); recipients were subsequently sacrificed the next day and spleen samples were assayed for CD45.2 precursor frequency counting. Data from single run are shown.

**b-c**, Frequencies of Pre5 and Pre7 germline precursors after 5K, 50K or 200K CD45.2 B cells were transferred. (**b**) Pre5; (**c**) Pre7. Each symbol represents a different mouse, n=5 or 6.

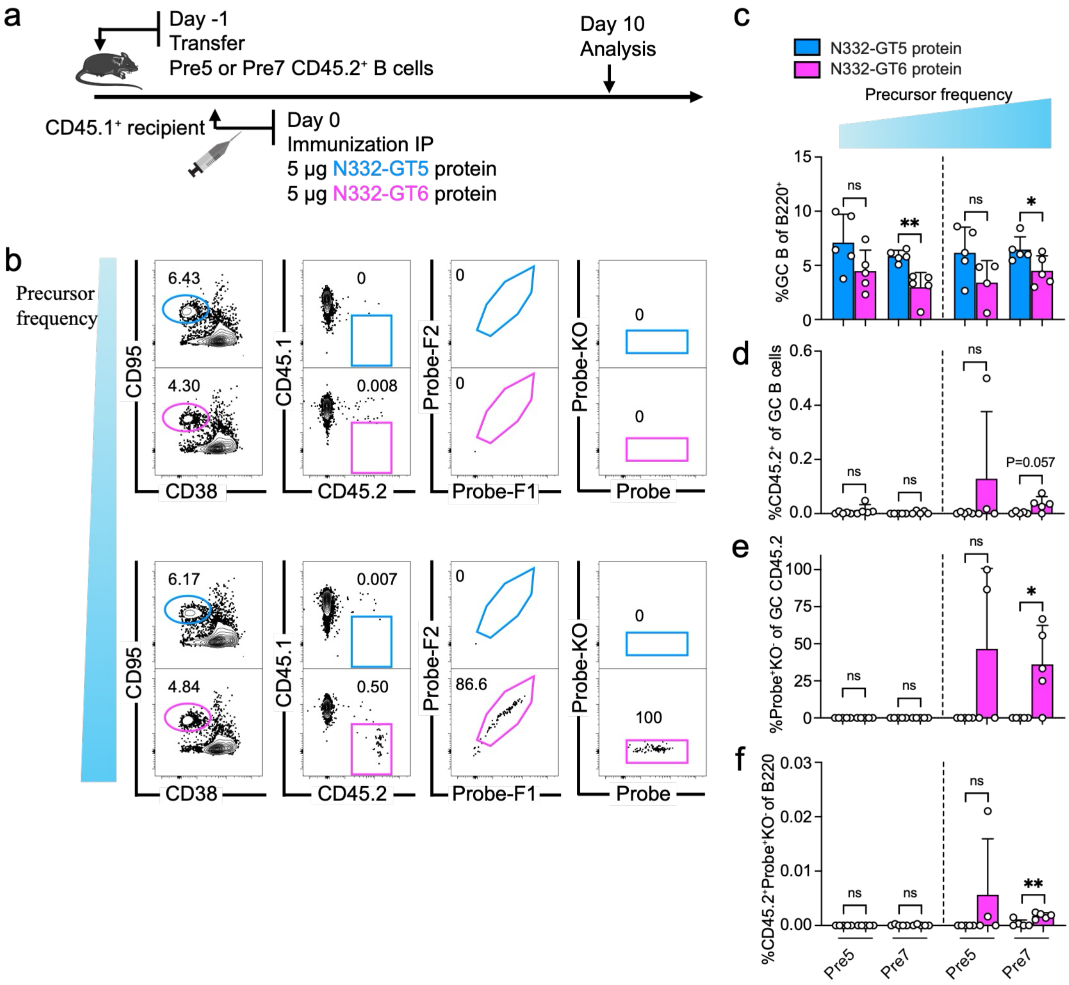


**Supplementary Fig. 5**

**Supplementary Fig. 5: Pre5 and Pre7 mouse model priming with N332-GT5 and N332-GT6 protein**

**a**, Schematic showing naïve CD45.2 pre5 and pre7 B cells at intermediate or high precursor frequencies adoptively transferred into CD45.1 WT mice (day -1). Recipients were subsequently immunized with either N332-GT5 or GT6 trimer protein formulated with Sigma adjuvant via intraperitoneal (IP) injection. Samples were assayed 10 dpi. Experiment with cells at high frequency was performed for twice, and data from single run are shown.

**b**, Gating strategy plots showing GC, CD45.2 cells in GC, antigen probe (N332-GT5, N332-GT6.2 or N332-GT6.3) binders of CD45.2 cells in GC and binding specificity after immunization using Pypre5 model.

**c-f**, Frequencies of GC B cells, $CD45.2^{+}$ B cell in GC, V3-glycan epitope specific ($N332\text{-}GT5^{+}KO^{-}$, $N332\text{-}GT6.2^{+}KO^{-}$ or $N332\text{-}GT6.3^{+}KO^{-}$) binders of GC $CD45.2^{+}$ cells gated as in (**b**) and CD45.2 epitope-specific binders in total B cells 10 dpi. Intermediate and high precursor frequency were broken by dashed line; each symbol represents a different mouse, n=5 in each condition except n=4 in Pypre5 high frequency primed with GT6. One-Way ANOVA analysis with Tukey's multiple comparison test or Brown-Forsythe and Welch ANOVA with Dunnett's T3 multiple comparison was applied depending on equality of variance. Bars indicate mean + SD. Not significant (ns); $*P < 0.05$; $**P < 0.01$.

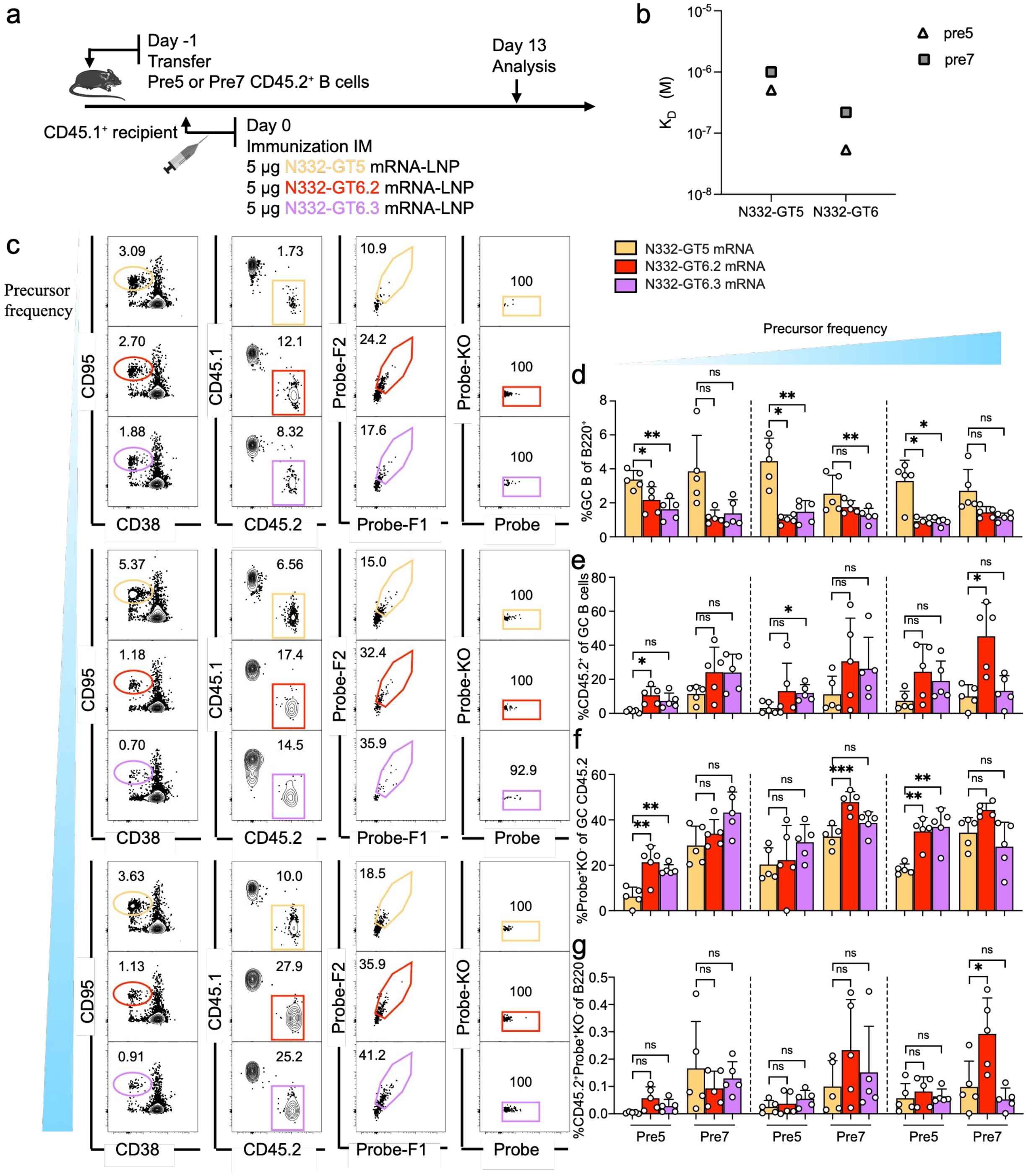


**Supplementary Fig. 6**

**Supplementary Fig. 6: Pre5 and Pre7 mouse model priming with N332-GT5, N332-GT6.2 and N332-GT6.3 mRNA-LNP**

**a**, Schematic showing naïve CD45.2 pre5 and pre7 B cells at low, intermediate or high precursor frequency adoptively transferred into CD45.1 WT mice (day -1); recipients were subsequently immunized with single N332-GT5, N332-GT6.2 or N332-GT6.3 mRNA-LNP via intramuscular (IM) injection. Samples were assayed 13 days post-immunization (dpi). Experiment with cells at high frequency was performed for at least twice, and data from single run are shown.

**b**, SPR affinity ($K_D$) for N332-GT5 and N332-GT6 binding to pre5 and pre7.

**c**, Gating strategy plots showing GC B cells, CD45.2 cells in GCs, antigen probe (N332-GT5, N332-GT6.2 or N332-GT6.3) binders among CD45.2 cells in GCs, and binding specificity after immunization. Representative flow cytometry panels from the Pypre5 model.

**d-g**, Frequencies of GC B cells, CD45.2$^+$ B cell in GC, V3-glycan epitope specific (N332-GT5$^+$KO$^-$, N332-GT6.2$^+$KO$^-$, or N332-GT6.3$^+$KO$^-$) binders among GC CD45.2$^+$ cells gated as in (**b**) and CD45.2 epitope-specific binders in total B cells 13 dpi. Low, intermediate and high precursor frequency were broken by dashed line; each symbol represents a different mouse, n=5 in each condition. One-Way ANOVA analysis with Tukey's multiple comparison test or Brown-Forsythe and Welch ANOVA with Dunnett's T3 multiple comparison was applied depending on equality of variance. Bars indicate mean + SD. Not significant (ns); *$P$ < 0.05; **$P$ < 0.01; ***$P$ < 0.001.

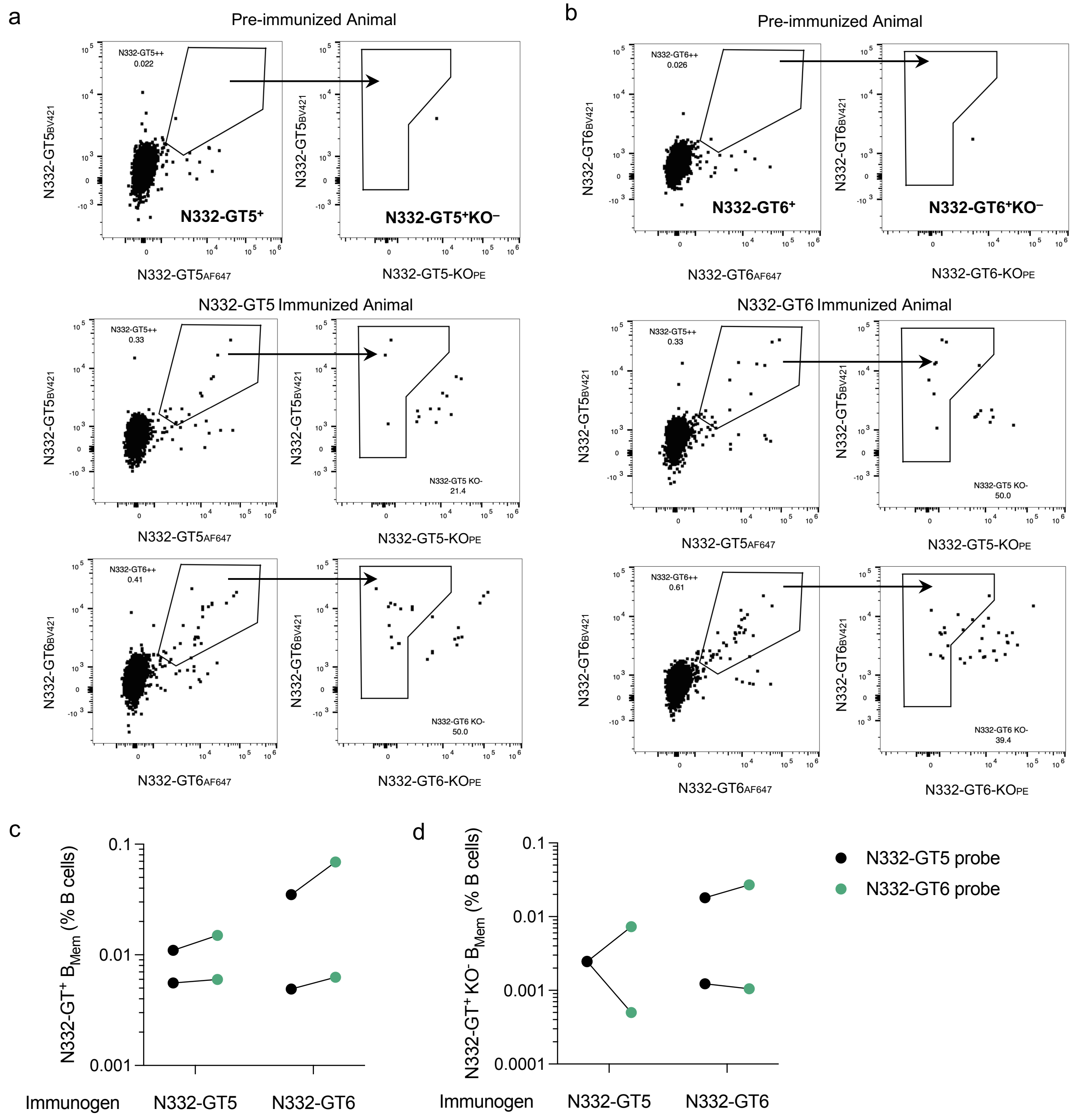


**Supplementary Fig. 7**

**Supplementary Fig. 7: Comparison of N332-GT5 and N332-GT6 probe staining in NHPs.**

**a,** Representative flow plots of N332-GT6-binding B cells prior to mRNA immunization (top). Representative flow plots comparing N332-GT5 and N332-GT6 probe binding across animals immunized with N332-GT5. PBMCs were stained from animals 2-weeks post-second dose of N332-GT mRNA (week 10; bottom).

**b,** Representative flow plots of N332-GT6-binding B cells prior to mRNA immunization (top). Representative flow plots comparing N332-GT5 and N332-GT6 probe binding across animals immunized with N332-GT6. PBMCs were stained from animals 2-weeks post-second dose of N332-GT mRNA (week 10; bottom).

**c,** Comparison of N332-GT5 and N332-GT6 WT probe binding among animals either immunized with N332-GT5 or N332-GT6 mRNA.

**d,** Comparison of N332-GT5 and N332-GT6 epitope-specific ($KO^{-}$) probe binding among animals either immunized with N332-GT5 or N332-GT6 mRNA.

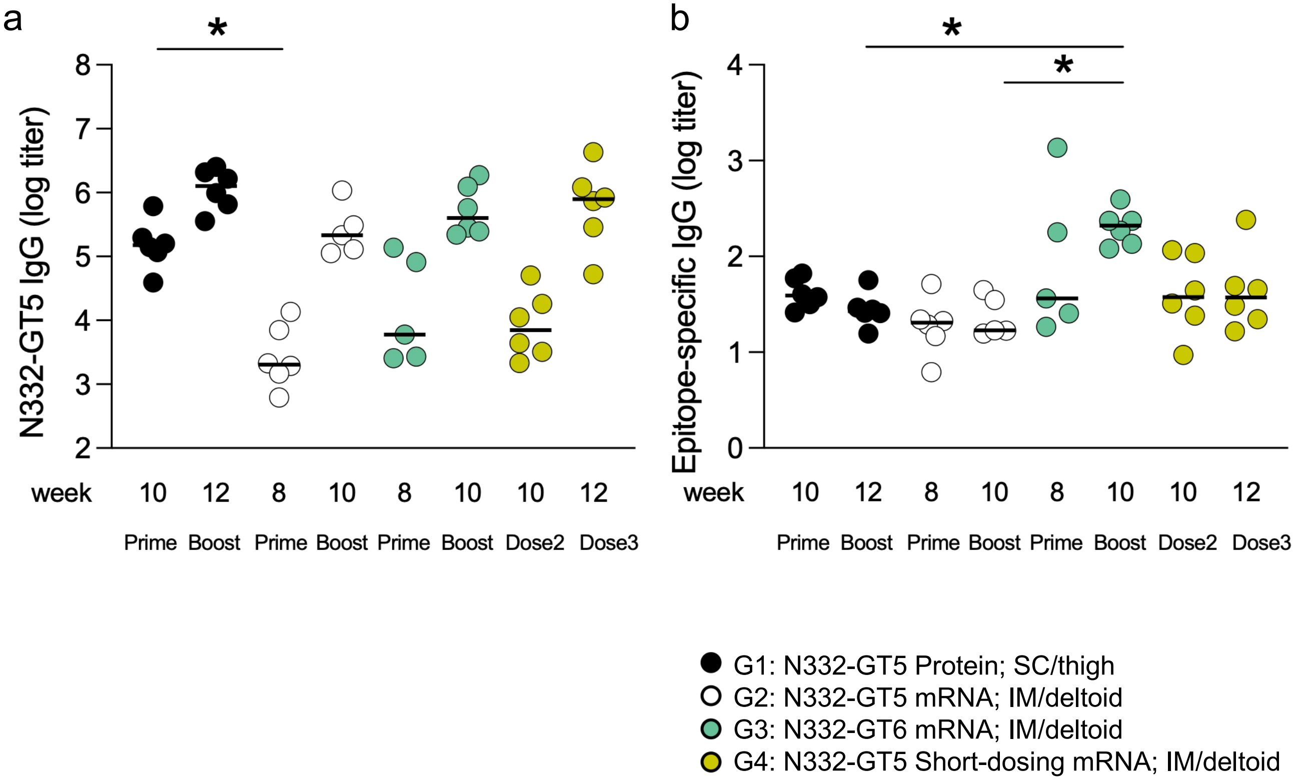


**Supplementary Fig. 8**

**Supplementary Fig. 8: ELISA serum responses to mRNA regimens in NHPs**

**a,** N332-GT5 IgG endpoint titers and **b**, epitope-specific titers 2-weeks post-prime or post-boost, as determined by ELISA. Bars indicate the medians. Significance was measured by Kruskal-Wallis test followed by Dunn's multiple comparisons test with adjustment for multiple comparisons. *$P < 0.05$, comparisons not marked were not significant.

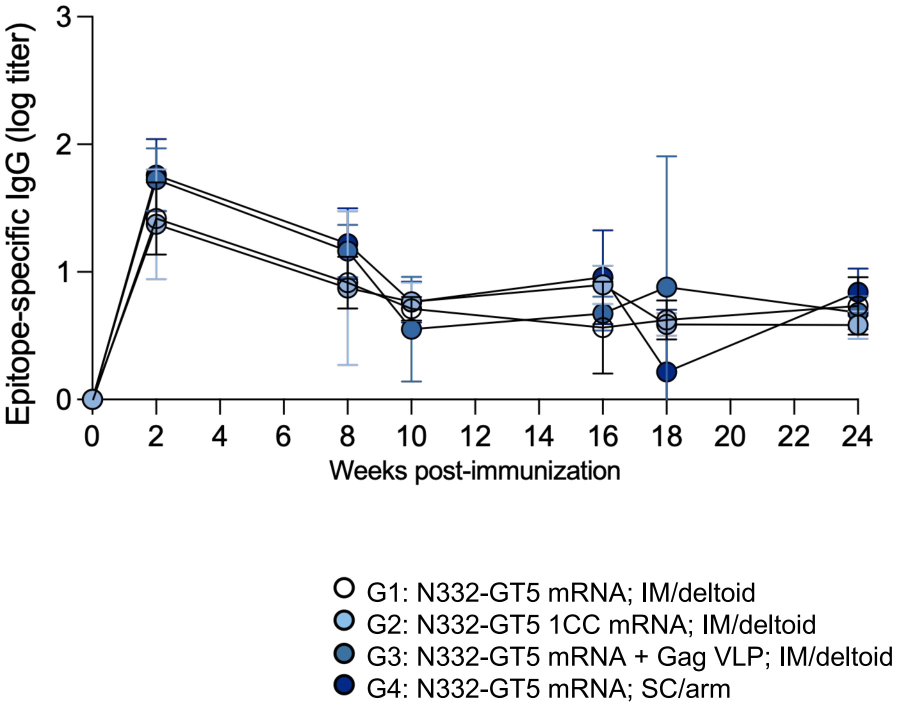


**Supplementary Fig. 9**

**Supplementary Fig. 9: Epitope-specific serum ELISA.**

Full timecourse of epitope-specific IgG endpoint titers as measured by ELISA. Geometric mean and SD are shown.

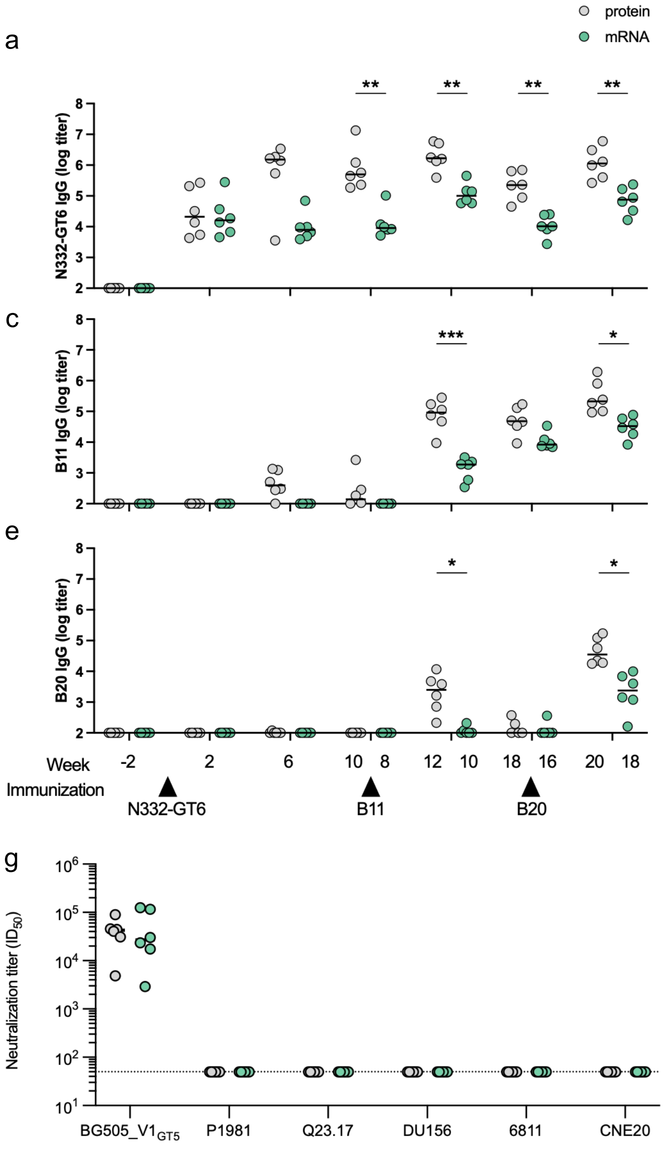


**Supplementary Fig. 10**

**Supplementary Fig. 10: Antibody responses and serum neutralization after heterologous boosting**

**a-f,** Full time course of IgG (**a,c,e**) endpoint titers and (**b,d,f**) epitope-specific titers to (**a,b**) N332-GT6, (**c,d**) B11, and (**e,f**) B20 measured by serum ELISA.

**g,** Serum from animals post-B20 booster immunization were tested for neutralization activity against a panel of wild-type HIV pseudoviruses and a variant of BG505 containing the N332-GT5 V1 loop, BG505_V1$_{GT5}$. BG18 is a positive control and Den3 is a negative control. Dotted line indicates the limit of detection (<50). Significance was measured using two-way ANOVA test with Šídák-adjusted P values for between-group comparisons at each timepoint. *P < 0.05, **P < 0.01, ***P < 0.001; ns, not significant.

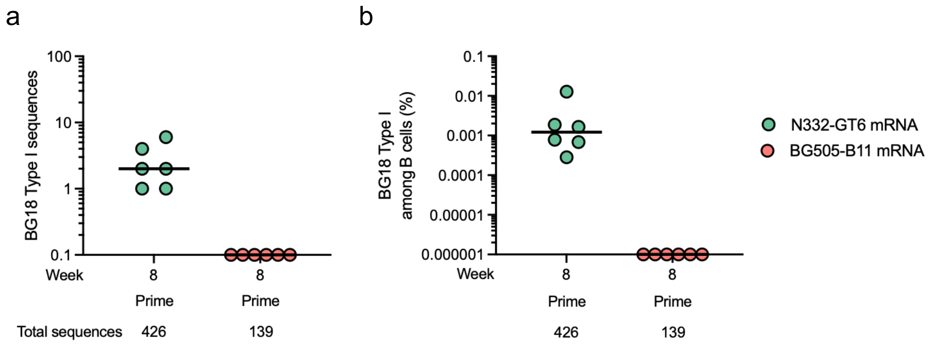


**Supplementary Fig. 11**

**Supplementary Fig. 11: Administration of B11 mRNA alone does not induce BG18 type I responses.**

**a,** Number of BG18 type I sequences identified post-priming immunization with N332-GT6 mRNA (NHP Study 1) or B11 mRNA (NHP Study 4).

**b,** Percentage of BG18 type I sequences among B cells.

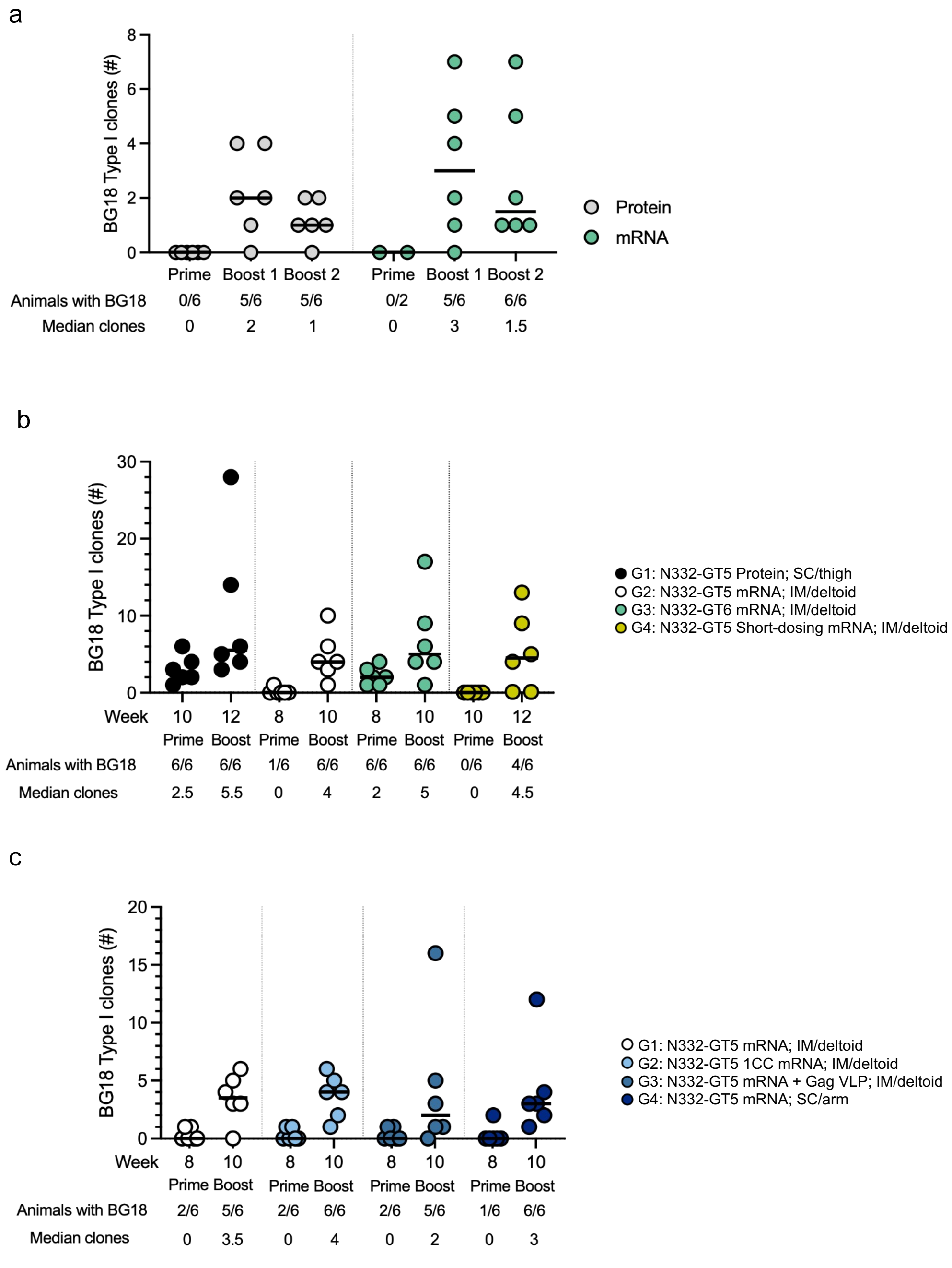


**Supplementary Fig. 12**

**Supplementary Fig. 12: Number of BG18-class precursor clonal lineages.**

**a,** Number of BG18 type I clones (distinct clonal lineages) recovered after priming and boosting with heterologous boosters in NHP Study 3.

**b,** Number of BG18 type I clones (distinct clonal lineages) recovered after priming and boosting with homologous N332-GT5 or N332-GT6 in NHP Study 1.

**c,** Number of BG18 type I clones (distinct clonal lineages) recovered after priming and boosting with homologous N332-GT5 mRNA constructs in NHP Study 2.

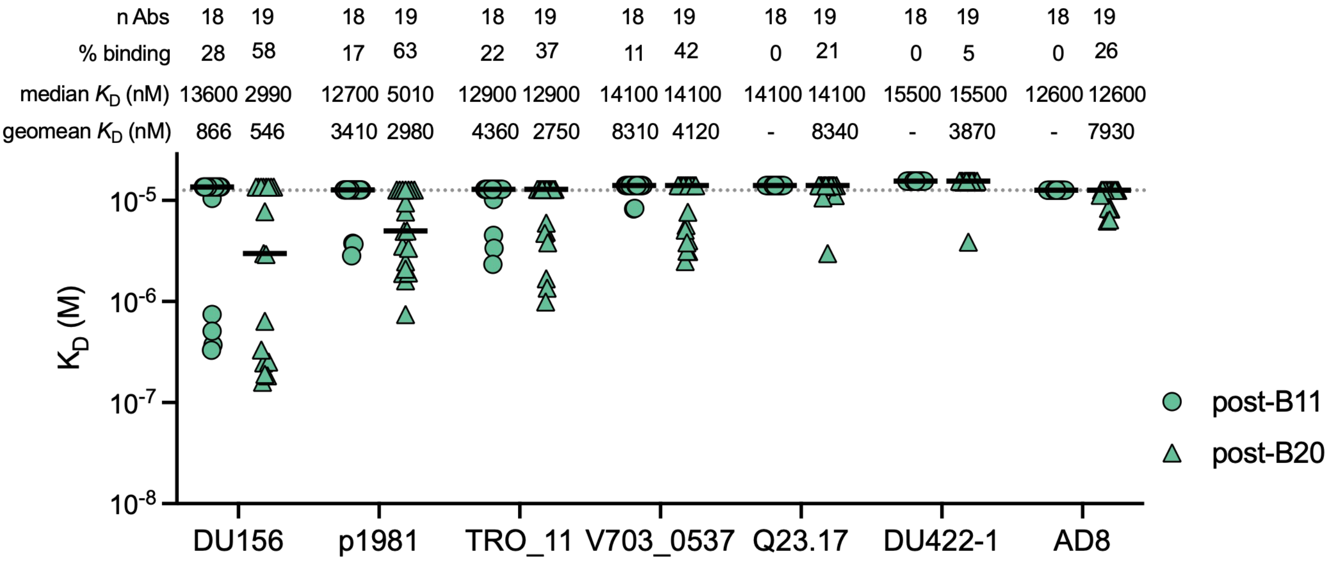


**Supplementary Fig. 13**

**Supplementary Fig. 13: Binding affinities of BG18 type I antibodies for wild-type trimers.**

SPR $K_D$s of BG18 type I antibodies isolated 2-weeks post-B11 and post-B20 booster immunization for wild-type trimers DU156, p1981, TRO_11, V703_0537, Q23.1, DU422-1, and AD8. Dotted line represents the limit of detection and thick lines represent median values.

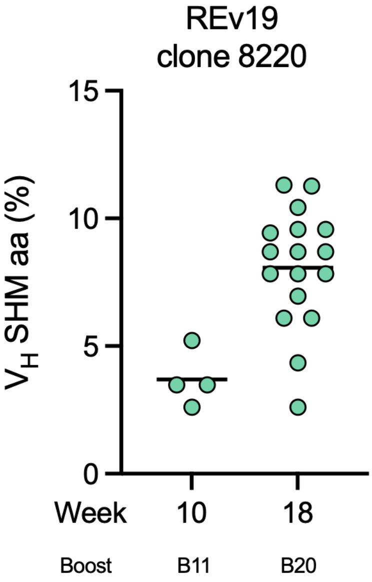


**Supplementary Fig. 14**

**Supplementary Fig. 14: Low somatic hypermutation in BG18-class lineage from animal Rev19**. VH gene amino acid SHM in the BG18 type I lineage of animal Rev19.
