## Supplemental Tables for "HIV broadly neutralizing antibody precursor maturation induced by mRNA vaccination in nonhuman primates"

**Supplementary table 1.** Flow Panel for sorting of N332-GT5 or N332-GT6-binding $B_{Mem}$ cells in NHPs

| Antibodies | Clone | Source | Cat # | Dilution |
|---|---|---|---|---|
| Streptavidin AF647 (N332-GT) | - | BioLegend | 405237 | - |
| TotalSeq-C Streptavidin BV421 (N332-GT) | - | BioLegend | custom | - |
| TotalSeq-C Streptavidin PE (N332-GT-KO) | - | BioLegend | custom | - |
| Streptavidin PE-Cy7 (B46) | - | BioLegend | 405206 | - |
| Streptavidin BV711 (B46) | - | BioLegend | 405241 | - |
| Streptavidin AF647 (B46-KO) | - | BD Biosciences | 613013 | - |
| Fixable Viability Dye APC-eFluor780 | - | Thermo Fisher Scientific | 65-0865-14 | 1:2000 |
| Mouse anti-human CD8a BV510 | RPA-T8 | BD Biosciences | 563256 | 1:100 |
| Mouse anti-human CD16 BV510 | 3G8 | BioLegend | 302048 | 1:100 |
| Mouse anti-human CD14 BV510 | M5E2 | BioLegend | 301842 | 1:100 |
| Mouse anti-human CD3 BV510 | SP34-2 | BD Biosciences | 740187 | 1:100 |
| Mouse anti-human CD20 BUV395 | 2H7 | BD Biosciences | 563782 | 1:100 |
| Mouse anti-human IgG BV605 | G18-145 | BD Biosciences | 563246 | 1:100 |
| Goat anti-human IgD AF488 | polyclonal | SouthernBiotech | 2030-30 | 1:50 |

**Supplementary table 2.** Flow Panel for sorting of N332-GT6-binding or B11-binding $B_{Mem}$ cells in NHPs

| Antibody | Clone | Source | Catalog Number | Dilution |
|---|---|---|---|---|
| TotalSeq-C0977 APC Streptavidin | - | BioLegend | 405197 | - |
| TotalSeq-C5222 BV421 Streptavidin | - | BioLegend | Custom | - |
| TotalSeq-C0953 PE Streptavidin | - | BioLegend | 405265 | - |
| PE-Cy7 Streptavidin | - | BioLegend | 405206 | - |
| BV711 Streptavidin | - | BioLegend | 405241 | - |
| BUV615 Streptavidin | - | BD Biosciences | 613013 | - |
| eBioscience Fixable Viability Dye eFluor 506 | - | Invitrogen | 65-0866-14 | 1:500 |
| Mouse anti-human CD3 APC-Cy7 | SP34-2 | BD Biosciences | 557757 | 1:100 |
| Mouse anti-human CD14 APC-Cy7 | M5E2 | BioLegend | 301820 | 1:100 |
| Mouse anti-human CD16 APC-eFluor780 | eBioCB 16 | Thermo Fisher Scientific | 47-0168-42 | 1:100 |
| Mouse anti-human CD20 PerCP-Cy5.5 | 2H7 | BioLegend | 302326 | 1:50 |
| Mouse anti-human CD27 BV650 | O323 | BioLegend | 302828 | 1:50 |
| Goat anti-human IgD Alexa Fluor 488 | polyclon al | Southern Biotech | 2030-30 | 1:40 |
| Mouse anti-human IgG BV786 | G18-145 | BD Biosciences | 564230 | 1:40 |
| Mouse anti-human IgM BUV395 | G20-127 | BD Biosciences | 563903 | 1:40 |

**Supplementary table 3.** Flow Panel for sorting of B20-binding $B_{Mem}$ cells in NHPs

| Antibody | Clone | Source | Catalog Number | Dilution |
|---|---|---|---|---|
| TotalSeq-C0996 Alexa Flour 647 Streptavidin | - | BioLegend | Custom | - |
| TotalSeq-C5225 BV421 Streptavidin | - | BioLegend | Custom | - |
| TotalSeq-C5231 BV421 Streptavidin | - | BioLegend | Custom | - |
| TotalSeq-C0951 PE Streptavidin | - | BioLegend | 405261 | - |
| TotalSeq-C5078 PE Streptavidin | - | BioLegend | Custom | - |
| PE-Cy7 Streptavidin | - | BioLegend | 405206 | - |
| BV711 Streptavidin | - | BioLegend | 405241 | - |
| BUV615 Streptavidin | - | BD Biosciences | 613013 | - |
| eBioscience Fixable Viability Dye eFluor 780 | - | Invitrogen | 65-0865-14 | 1:1000 |
| Mouse anti-human CD3 BV510 | SP34-2 | BD Biosciences | 740187 / 569486 | 1:100 |
| Mouse anti-human CD8 BV510 | RPA-T8 | BD Biosciences | 563256 | 1:100 |
| Mouse anti-human CD14 BV510 | M5E2 | BioLegend | 301842 | 1:100 |
| Mouse anti-human CD16 BV510 | 3G8 | BioLegend | 302048 | 1:100 |
| Mouse anti-human CD20 BUV395 | 2H7 | BD Biosciences | 563781 | 1:100 |
| Goat anti-human IgD Alexa Fluor 488 | Polyclonal | Southern Biotech | 2030-30 | 1:50 |
| Mouse anti-human IgG BV605 | G18-145 | BD Biosciences | 563246 | 1:100 |

**Supplementary table 4.** Primer sequences used for Rhesus macaque scBCR-seq library preparation

| | Target region | PCR | Sequence |
|---|---|---|---|
| V2_10X_forward_Bmix1 | - | Outer/Inner | GATCTACACTCTTTCCCTACACGACGC |
| IGHA_outer | IGHA | Outer | GGCTGGGATTTGTGTAGTGC |
| IGHM_outer | IGHM | Outer | CTCTCAGGACTGATGGGAAGC |
| IGHG_outer | IGHG | Outer | TTGTCCACCTTGGTGTTGCT |
| IGLC_outer | IGLC | Outer | GTCTCCACTCCCGCGTTGAC |
| IGKC_outer | IGKC | Outer | AGGTGTTGTCCTTGCTGTCC |
| IGHA_inner | IGHA | Inner | TCACGTTGAGTGGCTCCT |
| IGHM_inner | IGHM | Inner | AACGGCCACTTCGTTTGT |
| IGHG_inner | IGHG | Inner | AGCCCTGAGGACTGTAGGA |
| IGLC_inner | IGLC | Inner | ATCTGCCTTCCAGGCCA |
| IGKC_inner | IGKC | Inner | ACCTTCCACTTTACGCT |

**Supplementary table 5.** Cryo-EM data collection, refinement and validation statistics

| | MD65 N332-GT6 SOSIP + RM20A3 (EMD-79016) (PDB 38OQ) | SF162P3-B20 SOSIP + Rev19_06B + 3BNC117 (EMD-79015) (PDB 38OP) |
|---|---|---|
| **Data collection and processing** | | |
| Microscope | TFS Glacios 2 | TFS Glacios |
| Voltage (keV) | 200 | 200 |
| Camera | TFS Falcon 4i | TFS Falcon 4 |
| Collection mode | Counting | Counting |
| Magnification | 190,000x | 190,000x |
| Pixel size at detector (Å) | 0.718 | 0.725 |
| Total electron exposure (e–/Å$^2$) | 49.9 | 45.0 |
| Exposure rate (e-/pixel/sec) | 12.3 | 6.15 |
| Number of EER frames | 40 | 40 |
| Defocus range (μm) | -0.8 to -1.7 | -0.8 to -2.0 |
| Automation software | EPU | EPU |
| Micrographs collected (no.) | 5,001 | 4,674 |
| Micrographs used (no.) | 4,749 | 2,251 |
| Initial particle images (no.) | 585,264 | 279,698 |
| Final particle images (no.) | 54,181 | 120,115 |
| Symmetry | C3 | C1 |
| Final map pixel size | 1.034 | 0.932 |
| Map resolution (masked/unmasked Å) | 3.3/3.9 | 3.7/4.3 |
| FSC threshold | 0.143 | 0.143 |
| Map sharpening *B* factor (Å$^2$) | -86 | -90 |
| Map resolution range (Å) | 2.5-4.5 | 3.0-5.5 |
| **Refinement** | | |
| Initial model used (PDB code) | 8T4K | AlphaFold 3 |
| Refinement package | Phenix RSR | Phenix RSR |
| Model resolution (Å) | 3.4 | 4.0 |
| FSC threshold | 0.5 | 0.5 |
| EMRinger score | 3.00 | 2.03 |
| CC (mask) | 0.87 | 0.81 |
| *Model composition* | | |
| Non-hydrogen atoms | 19,563 | 21,344 |
| Protein residues | 2,373 | 2,560 |
| Ligands | NAG: 72 | NAG: 66 |
| | BMA: 3 | BMA: 8 |
| | MAN: 3 | MAN: 7 |
| *Mean B factors (Å$^2$)* | | |
| Protein | 74 | 151 |
| Ligand | 99 | 157 |
| *R.m.s. deviations* | | |
| Bond lengths (Å) | 0.006 | 0.004 |
| Bond angles (°) | 0.776 | 0.754 |
| *Validation* | | |
| MolProbity score | 1.24 | 1.08 |
| Clashscore | 0.70 | 0.31 |
| Poor rotamers (%) | 0.54 | 0.22 |
| *Ramachandran plot* | | |
| Favored (%) | 91.27 | 92.51 |
| Allowed (%) | 8.73 | 7.49 |
| Disallowed (%) | 0.00 | 0.00 |
| Cβ outliers (%) | 0.00 | 0.00 |
| CaBLAM outliers (%) | 4.13 | 5.13 |
